# A transcriptomic axis aligns with in vivo functional dynamics in hippocampal inhibitory circuits

**DOI:** 10.64898/2026.04.07.716935

**Authors:** Hyun Choong Yong, Stephanie A. Herrlinger, Margaret E. Conde Paredes, Cliodhna K. O’Toole, Juyoun Yoo, Bovey Y. Rao, Tiberiu S. Mihaila, Jingcheng Shi, Subhrajit Dey, Erdem Varol, Attila Losonczy

**Affiliations:** Mortimer B. Zuckerman Mind Brain Behavior Institute, Columbia University; New York, NY, USA; Doctoral Program in Neurobiology and Behavior, Columbia University; New York, NY, USA; Tandon School of Engineering, New York University; New York, NY, USA; Peter O’Donnell Jr. Brain Institute, University of Texas Southwestern Medical Center; Dallas, TX, USA; Department of Neuroscience, University of Texas Southwestern Medical Center; Dallas, TX, USA

## Abstract

Linking molecular identity to function *in vivo* at single-cell resolution remains an outstanding challenge in neuroscience. Here, we bridge this gap in the mouse hippocampus with an end-to-end pipeline of cell-resolved two-photon imaging and spatial transcriptomics. CA1 interneurons exhibiting heterogeneous physiological responses during a virtual-reality navigation task were *post hoc* clustered by gene expression into 5 GABAergic subclasses and 14 types. Physiological responses of individual cells aligned with a transcriptomic axis, and a classifier trained on physiological features alone recovered the same ordered organization. Our approach establishes a direct, scalable framework for linking *in vivo* circuit dynamics to constituent cell identity, revealing a transcriptomic axis that encompasses the structural and functional diversity of hippocampal inhibitory neurons.

**One-Sentence Summary:** Tracking neurons from behavior to spatial transcriptomics links *in vivo* function to molecular identity in the hippocampus.

## Main Text

A long-standing unresolved question in neuroscience is how the molecular diversity of mammalian cell types relates to their functional roles *in vivo.* Mammalian cortical circuits comprise hundreds of transcriptomically distinct cell types (*1–12*), yet the extent to which this diversity maps onto the functional heterogeneity observed *in vivo* remains elusive. Traditional approaches have relied on (1) immunohistochemistry to assign cell identity post hoc to *in vivo* recorded neurons (*13–21*), (2) transgenic lines to record from genetically restricted subpopulations of neurons (*22–25*), (3) activity-dependent markers to label functionally engaged neuronal subsets (*26–28*), or (4) advanced methods for isolating physiologically defined cell types (*29, 30*). While these strategies have yielded foundational insights, they do not provide a scalable, direct link between transcriptomic identity and *in vivo* physiological function across cell types at single-cell resolution.

Recent advances in spatial transcriptomics (ST) provide a potential solution to bridge these gaps by enabling high-throughput molecular identification of neurons within intact tissue at single-cell resolution (*31–34*). However, realizing this potential for neurons recorded *in vivo* requires robust integration across three distinct modalities: *in vivo* imaging, *ex vivo* imaging, and spatial transcriptomic measurements. Because these modalities differ in resolution, scale, and tissue state, linking *in vivo* circuit dynamics to high-resolution *post hoc* spatial transcriptomics remains a core technical barrier.

We focused on GABAergic interneurons (INs) in hippocampal area CA1 to establish an end-to-end pipeline for physiologically correlated spatial transcriptomics. CA1 INs are highly diverse at molecular, anatomical, and functional levels, and they play central roles in regulating CA1 pyramidal cell activity and network dynamics (*35–39*). The relationship between transcriptomic identity and *in vivo* physiological function, however, remains poorly defined. Moreover, it is unclear whether transcriptomic variation beyond canonical marker genes systematically correlates with the functional heterogeneity observed *in vivo*.

To address this, we combined *in vivo* two-photon Ca^2+^ imaging of CA1 INs in head-fixed mice performing a virtual reality spatial navigation task (VR) with *post hoc* MERSCOPE (MERFISH-based ST) of the same tissue (*31, 40*). Using a pan-neural 500-gene panel (see Methods) and a multimodal registration pipeline, we obtained matched physiological and transcriptomic profiles from 610 CA1 INs (n=4 animals). This approach enabled transcriptomics-based classification of imaged CA1 INs into five major GABAergic subclasses (*Pvalb*, *Sst*, *Lamp5*, *Vip*, and *Chrm2*) and fourteen types defined from transcriptomic similarity while preserving laminar information.

We found that physiological responses to a novel context were heterogeneous across CA1 INs, and that a subset of physiological features, related to velocity, activity level, and reward covaried with position along a transcriptomic axis derived solely from gene expression. This axis corresponded to the second transcriptomic principal component (tPC2), whose gene loadings spanned from *Pvalb*-associated markers to *Lamp5*-associated markers. As an orthogonal test, a classifier trained on physiological features alone showed that subclass confidence margins varied smoothly along tPC2 position, consistent with graded association between physiological feature space and position along this transcriptomic axis. Collectively, our results establish a multimodal framework for directly linking gene expression to *in vivo* physiology in hippocampal circuits and suggest a graded structure of CA1 IN functional heterogeneity along a transcriptomic axis.

### Registration of imaged hippocampal INs across imaging modalities

To identify imaged CA1 INs across imaging modalities and correlate gene expression with physiology, we first established a cross-modal registration pipeline. GABAergic INs were labeled by injecting a Cre-dependent GCaMP8s virus (AAV1-hSyn-FLEX-jGCaMP8s-WPRE, Methods) into the dorsal CA1 of VGAT-Cre mice. Mice were then implanted with an imaging cannula-window over the dorsal CA1 and trained to navigate a linear VR environment with fixed reward locations. After the mice were well-trained in this task, they were imaged during a context-switch paradigm in which familiar and novel VR contexts alternated in blocks.

To capture *in vivo* GCaMP-Ca^2+^ activity dynamics of INs located in distinct CA1 sublayers, we performed multi-plane two-photon imaging in two separate sessions, targeting the *stratum oriens* and *stratum pyramidale*, respectively (Fig. 1A). This approach enabled us to sample INs that differ in their anatomical and transcriptional identities (*36, 41–43*). After completing *in vivo* functional imaging, we acquired a high-resolution *in vivo* two-photon z-stack volume (Fig. 1B) to serve as a structural reference for cross-modal registration. We subsequently extracted the brain and sectioned the imaged area into 12-µm thick horizontal slices to match the orientation of the functional imaging planes. These sections were imaged *ex vivo* using confocal microscopy to capture endogenous GCaMP8s signals. Subsequently, we performed MERFISH with the MERSCOPE PanNeuro Cell Type Mouse Panel (500-gene panel, Methods).

**Fig. 1.**
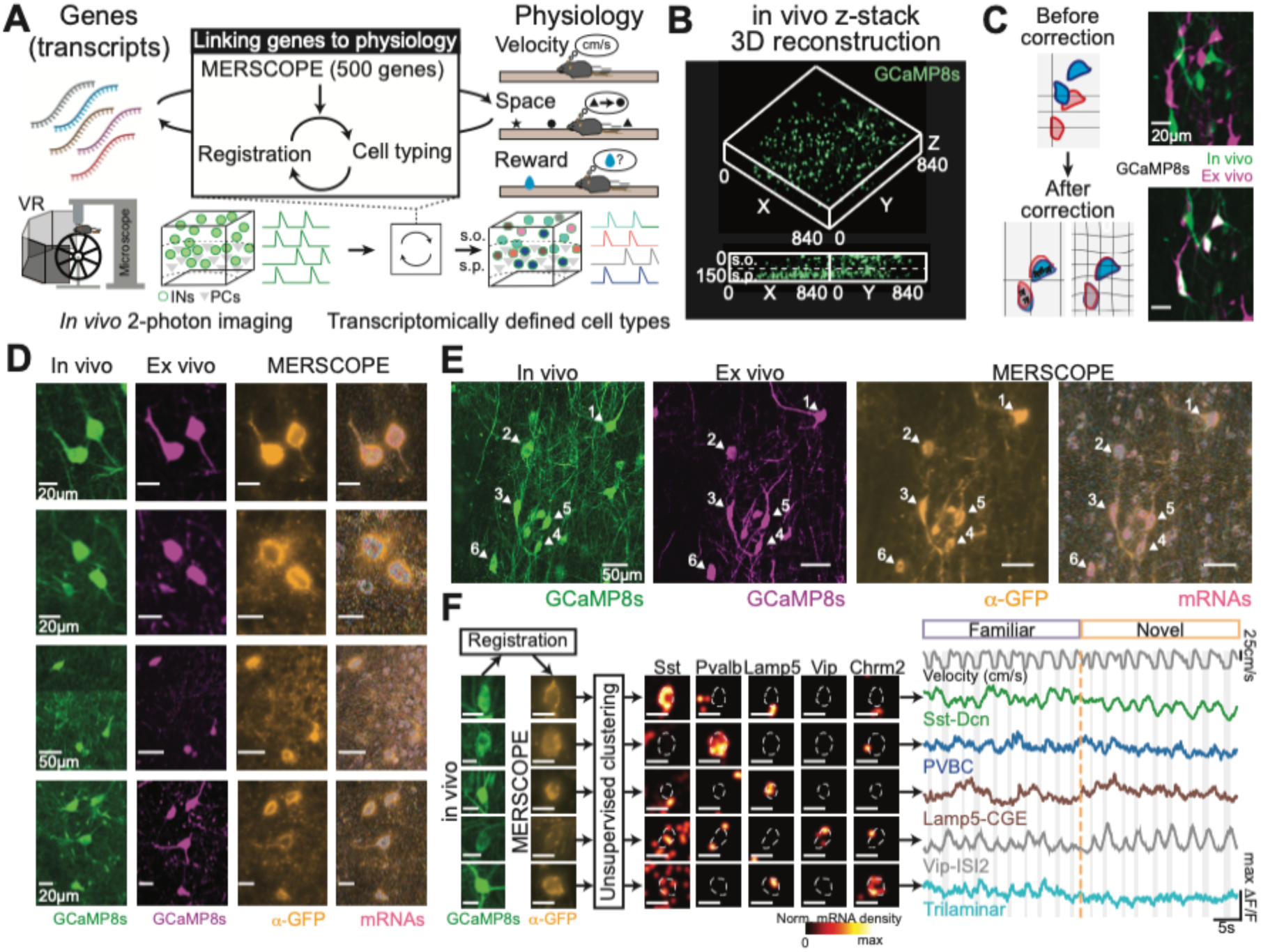
Identification of imaged hippocampal CA1 INs across imaging modalities. (**A**) Schematic of the experimental framework linking gene expression (transcripts) and *in vivo* physiology. Hippocampal CA1 interneurons (INs) expressing GCaMP8s were imaged *in vivo* to capture their physiological responses to behavioral correlates. Subsequently, MERSCOPE was performed with a predesigned 500-gene panel on the same brain tissue. Registration between *in vivo*, *ex vivo*, and MERSCOPE enabled identification of the same cells across modalities, followed by unsupervised clustering to assign transcriptomic cell types. Refer to *fig. S1* for the detailed workflow. (**B**) Example 3D reconstruction of the *in vivo* z-stack showing GCaMP8s-expressing CA1 INs. Bottom is the orthogonal view of the reconstructed stack. (**C**) Schematic of the registration pipeline. Before correction, INs are misaligned between *in vivo* (green) and *ex vivo* (magenta) images. After applying deformations and transformations (see Methods and *fig. S2*), images are aligned across modalities, allowing the identification of the same cells between *in vivo* and *ex vivo* images. (**D**) Examples of individual INs registered across modalities: *in vivo* GCaMP8s (green), *ex vivo* GCaMP8s (magenta), and anti-GFP antibody staining from MERSCOPE (yellow). Individual transcripts detected with MERSCOPE are shown as overlaid scatters (right, pink) (**E**) Example field of view (FOV) registered across modalities, demonstrating the scalability of our registration pipeline. (**F**) Examples of registered cells (left) with gene expression of canonical marker genes (middle; *Sst*, *Pvalb*, *Lamp5*, *Vip*, and *Chrm2* selected from the predesigned 500 gene panel) and segmentation outlines (gray dashed lines). Transcript density heatmaps were max-normalized to highlight the relative locations of detected mRNAs. Calcium traces colored by their cell type found through unsupervised clustering (right). This experimental framework successfully combines *in vivo* 2-photon imaging, behavior and MERSCOPE to identify cell types and link gene expression with physiology. For (**F**), the scale bar is 20-µm. Refer to *movie S1* for an additional visualization of the registration process.

Using GCaMP8s expression in INs as sparse fiducial markers, we manually registered the *ex vivo* images to the *in vivo* z-stacks. Maximal correspondence was achieved through a series of transformations and deformations with our established method (*44*) (Fig. 1C, fig. S1 and movie S1; Methods). This procedure produced a robust alignment of *in vivo* and *ex vivo* images, as supported by significant increases in normalized cross-correlation and mutual information metrics (fig. S1, B and C). Because the tissue-clearing step of the MERSCOPE protocol degrades GCaMP fluorescence, we incorporated anti-GFP immunolabeling as an additional visual reference for GCaMP-expressing cells in MERSCOPE image (orange, Fig. 1, D and E). Subsequently, we used DAPI staining applied at the beginning of each MERSCOPE run to register nuclei between the *ex vivo* and MERSCOPE images and identified the same nuclei across the two modalities (fig. S2). Individual cells were matched based on the Euclidean distance between segmented masks from *ex vivo* and MERSCOPE images (fig. S1, A, D). Segmented *in vivo*, *ex vivo*, and MERSCOPE GCaMP+ cells were then manually confirmed as matched cells in their registered 3D space. We applied this pipeline to 4 mice, retrieving a total of 610 INs (152.5 ± 24.5 cells per mouse, mean ± S.D.). Together, this registration pipeline enables reliable tracking of individual INs across *in vivo*, *ex vivo,* and MERSCOPE imaging modalities, thereby enabling direct linkage of *in vivo* physiological properties to gene expression profiles.

### Classification of INs based on transcriptomic profiles

To examine the transcriptomic profiles of the imaged cells, we performed unsupervised clustering and marker-gene-based classification on cells from all imaged tissues (fig. S3). Cells were segmented based on DAPI and cell boundary signals (fig. S3, top, Methods). After MERSCOPE images were collected from all sections, we manually segmented out non-hippocampal tissues (fig. S3, middle top). Spatial transcriptomes of a final total of 274K hippocampal cells were obtained from MERSCOPE images after applying quality control measures (fig. S3, middle bottom, Ext. Data fig. 1, A-G). We then applied unsupervised Leiden clustering and classified the clusters into 5 major cell classes (CA1 Pyramidal Neurons, GABAergic INs, Oligodendrocytes & Oligodendrocyte Progenitor Cells, Astrocytes, and Vascular Cells), based on differential expression of established cell-type marker genes (Fig. 2A, Ext. Data fig. 1, H-K; see Methods).

**Fig. 2.**
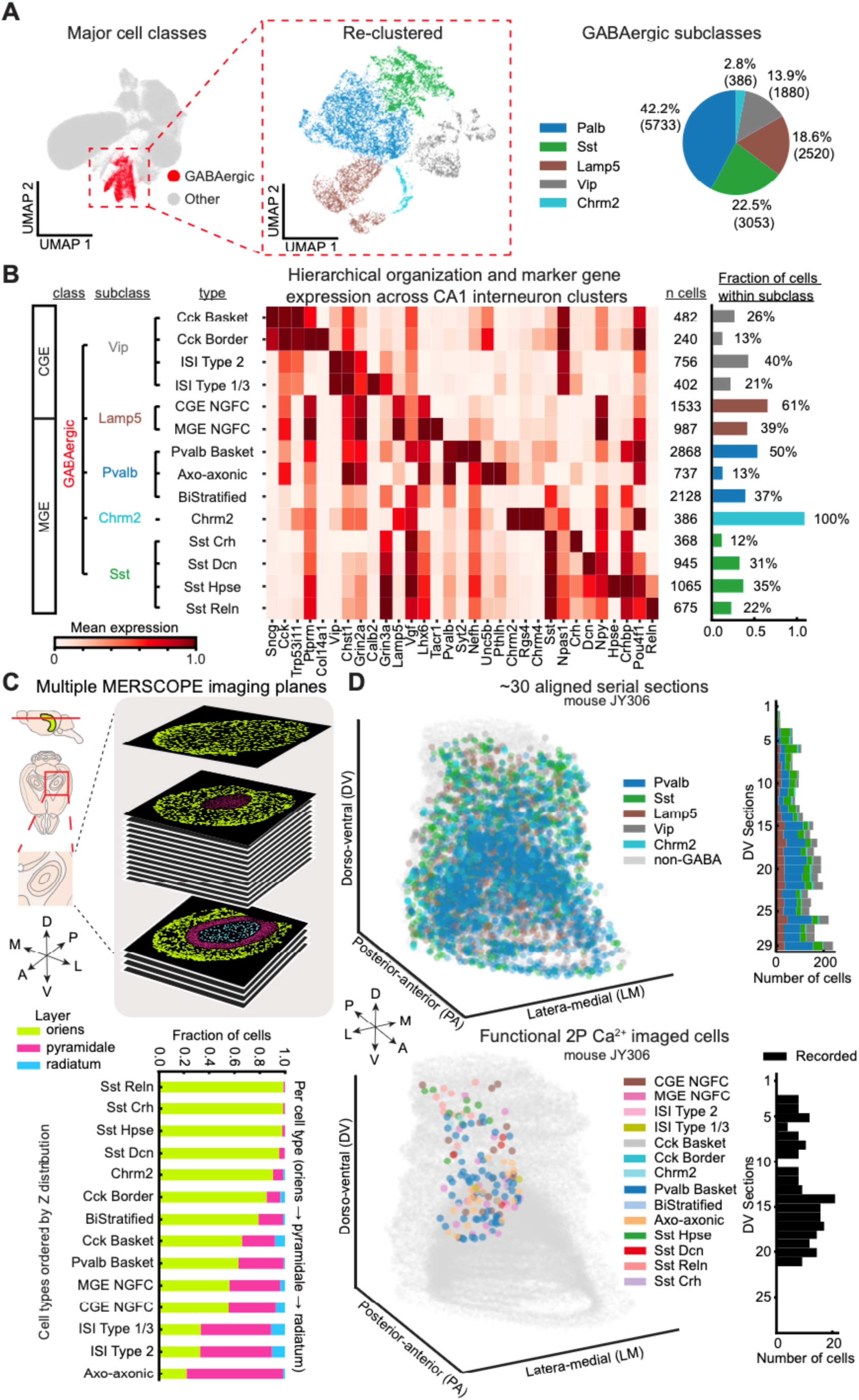
Spatial transcriptomics-based molecular classification revealing 3D laminar organization of 14 GABAergic interneuron types in hippocampal CA1. (**A**) UMAP of major cell classes (left) and GABAergic subclasses (middle) identified by MERSCOPE. Pie chart shows proportions of GABAergic subclasses (right). n=13,572 GABAergic cells. (**B**) Hierarchical classification of GABAergic interneurons from all cells from all animals collected, showing class, subclass, and type levels. Heatmap shows marker gene expression across 14 cell types. Numbers indicate cell counts; bars show the fraction within each subclass. (**C**) Laminar distribution of interneuron types across CA1 hippocampal layers. Top: Schematic shows multiple planes imaged with MERSCOPE. Bottom: fraction of each cell type located in *stratum oriens*, *pyramidale* or *radiatum*. Data from all mice. (**D**) Example 3D spatial reconstruction from ∼30 aligned serial sections (mouse JY306). Top: cells are color-coded by subclass. Bottom: same cells as top but showing functionally imaged cells (*in vivo* 2-photon imaging) color-coded by cell type. Histogram shows recorded cell counts by depth (DV, dorsoventral).

The GABAergic IN class was re-clustered to identify 5 major subclasses, defined by canonical IN subclass markers: *Pvalb*, *Sst*, *Lamp5*, *Vip*, and *Chrm2* (Fig. 2A, Ext. Data fig. 1, L-N). We mapped our cells to a reference MERFISH dataset (*2*) (see Methods) to compare our class and subclass level annotations with the hierarchical taxonomy from the Allen Brain Cell Atlas and found that they aligned with GABAergic IN subclasses from the reference dataset (fig. S4, A and B).

The reclustering of the GABAergic IN class revealed five major subclasses: parvalbumin-expressing (*Pvalb*; 42.2%, n=5,733 cells), somatostatin-expressing (*Sst*; 22.5%, n=3,053 cells), *Lamp5*-expressing (*Lamp5*; 18.6%, n=2,520 cells), vasoactive intestinal peptide-expressing (*Vip*; 13.9%, n=1,880 cells), and trilaminar/long-projecting *Chrm2*-expressing neurons (*Chrm2*; 2.8%, n=386 cells) (Fig. 2A, Ext. Data fig. 1, L-N). *Pvalb* and *Sst* INs together comprise 64.7% of the CA1 INs (n=8,786) (Fig. 2A). Overall subclass proportions were concordant with previous single-cell RNA sequencing studies for mouse hippocampus (*1, 45*). *Lamp5-*expressing INs have been largely associated with caudal ganglionic eminence (CGE)-derived lineages characterized by *Id2* expression across the cortex, including neurogliaform cell (NGFC) types (*10*). The transcriptional profile of the *Chrm2* subclass closely matched that of retrohippocampal-projecting INs in the Allen Brain Cell Atlas (fig. S4B) (*1, 10*). We then further reclustered each subclass to resolve cell types, which can be segregated by developmental origin into CGE-derived and medial ganglionic eminence (MGE)-derived cell type populations (Fig. 2B) (*36, 46*). Each clustering step was determined based on optimal silhouette and adjusted rand index scores, and relevant canonical type names were given with justification from the literature, resulting in 14 final cell types (Fig. 2B, Ext. Data fig. 1, O-KK, Methods).

Seeking to independently validate our classification, we tested whether well-established anatomical differences in interneuron position were recapitulated in our identified cells. Examination of the laminar distribution of each IN type across CA1 strata confirms cell types exhibited layer preferences consistent with their known morphologies and connectivity (Fig. 2C, fig. S5, A-C) (*16, 36, 42*). The *Pvalb* as well as the *Vip* subclasses localized predominantly to *stratum pyramidale* and superficial *stratum radiatum*, while the *Sst* and *Chrm2* subclasses concentrated in *stratum oriens*, patterns concordant with decades of anatomical characterization (*16, 35*). By aligning approximately 30 serial sections per mouse, we reconstructed a three-dimensional molecular map of CA1 INs spanning the dorsoventral, anteroposterior, and mediolateral axes (Fig. 2D, top). Together, these establish a transcriptomic and spatial taxonomy of CA1 INs, providing a multimodal reference for assigning molecular and spatial identities to CA1 INs imaged *in vivo*.

### Context-dependent in vivo activity dynamics of transcriptomically identified INs

After assigning transcriptomic labels to the functionally imaged CA1 INs, we investigated how physiological activity patterns in a reward-guided VR-context navigation task (Fig. 3, A and B) varied with transcriptomically defined subclasses and types. We first focused on velocity modulation and observed a robust relationship between velocity and GCaMP-Ca^2+^ activity (ΔF/F), as previously reported (*13, 14, 22, 23, 47, 48*) (Fig. 3D, fig. S6A). The magnitude of this modulation was context-dependent: the *Pvalb* subclass exhibited a significant reduction in their velocity modulation upon exposure to the novel context, driven largely by the *Pvalb* Basket and Axo-axonic (AAC) types, whereas the *Chrm2* subclass showed an increase (Fig. 3D, fig. S6B).

**Fig. 3.**
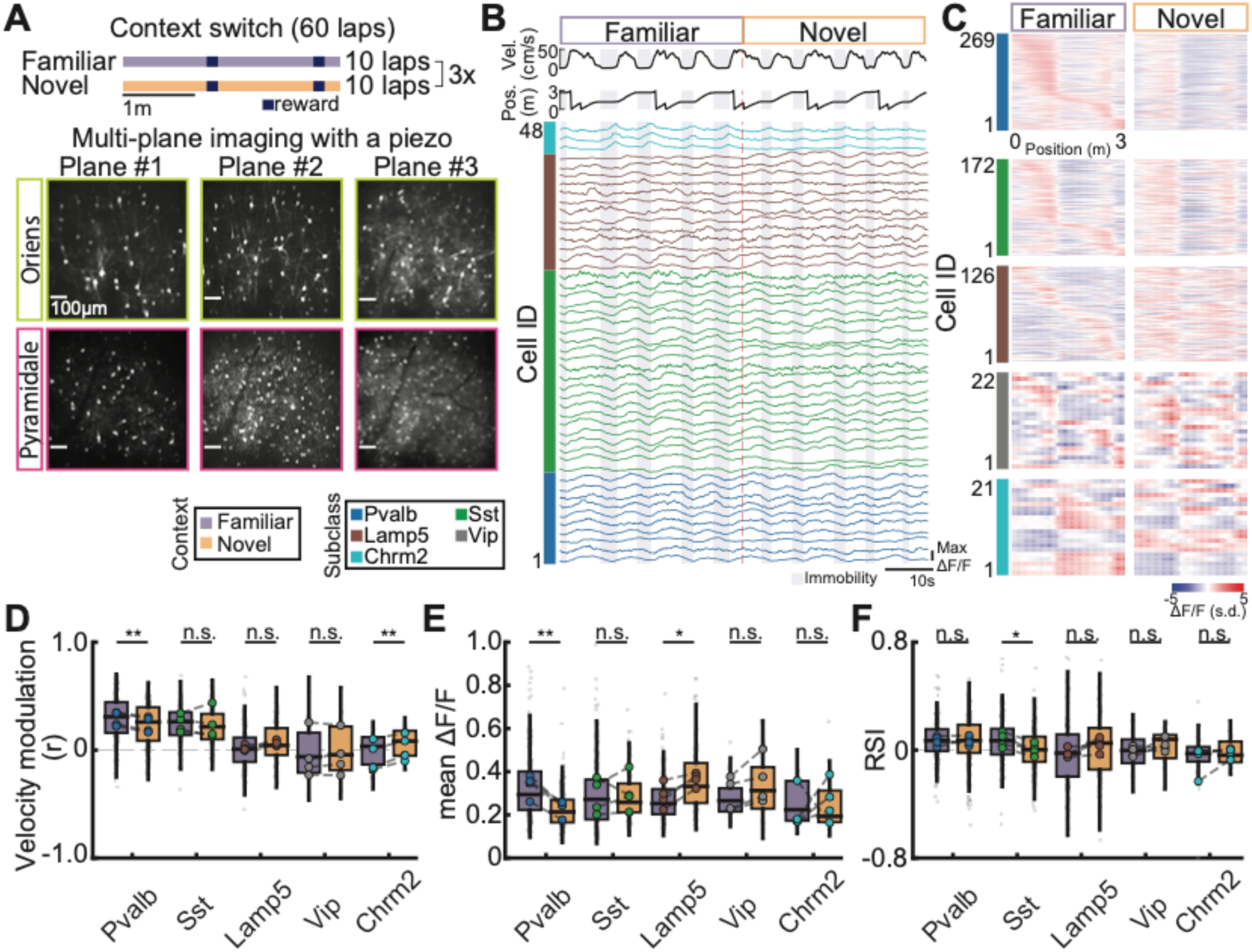
Heterogeneous context-dependent adaptations of transcriptomically identified CA1 INs. (**A**) Context-switch paradigm and example imaging field-of-views (FOVs) from *strata oriens* and *pyramidale* imaging sessions. Mice alternated between familiar and novel contexts (10 laps per context x 3 blocks). To maximize the number of CA1 INs imaged simultaneously, we imaged 3 planes per session, with each plane ∼30-µm apart. In both imaging sessions, mice were exposed to a novel environment that they had never seen before. (**B**) Example ΔF/F traces sorted by transcriptomically defined CA1 IN subclass imaged within the same session. (**C**) Rate maps, sorted by familiar context peak locations, for each CA1 IN subclass. Rate maps were z-scored using both familiar- and novel-context rate maps. (**D**) Velocity modulation grouped by each subclass. A significant difference was observed in *Pvalb* and *Chrm2* subclasses between familiar and novel contexts (*Pvalb*: t(3)=10.58, p=0.0018; *Chrm2*: t(3)=−6.57, p=0.0072; n=4 animals). (**E**) Changes in mean ΔF/F between novel and familiar contexts for each subclass. A significant change in mean ΔF/F for *Pvalb* and *Lamp5* subclasses (*Pvalb*: t(3)=9.63, p=0.0024; *Lamp5*: t(3)=−4.59, p=0.019; n=4 animals). (**F**) Changes in reward selectivity index (first vs. second reward, see Methods). Positive values indicate that ΔF/F is higher before the first reward zone compared to the second reward zone. *Sst* subclass shows a significant loss of reward preference in the novel context (*Sst*: t(3)=3.47, p=0.04; n=4 animals). For (**D**) through (**F**), boxplots indicate the distribution of individual cells (gray), and colored circles represent animal means.

We next asked whether *in vivo* activity levels and reward response differ across transcriptomically defined subclasses and types, independent of velocity. Because IN activity is strongly modulated by locomotion, we regressed out the velocity component from the calcium activity prior to analysis (see Methods) (*15, 49*). We then assessed context-dependent changes in mean activity (mean ΔF/F). While most subclasses maintained stable activity levels, the *Pvalb* subclass showed a significant reduction in the novel context, whereas the *Lamp5* subclass exhibited a significant increase (Fig. 3E). Finally, we examined reward selectivity, the preference for specific reward sites. The *Pvalb* subclass consistently exhibited a preference for the first reward zone in both contexts while the *Lamp5* subclass showed no significant bias (Fig. 3, C and F). In contrast, the *Sst* subclass displayed a significant context-dependent change with their preference for the first reward zone diminishing in the novel context (Fig. 3F). Together, these results demonstrate that the physiological response to novelty and reward is not uniform, but differs across transcriptomically defined subclasses and types.

### Organization of heterogeneous physiological responses along transcriptomic axes

Our analyses linking gene expression and functional dynamics revealed cell-type-dependent differences in CA1 INs between familiar and novel contexts (Fig. 3, fig. S6 and S7). These results raise the question of whether such heterogeneity reflects random variability or shows a relationship with any underlying transcriptomic structure. To address this question, we performed dimensionality reduction on the cell-by-gene matrix of the imaged and registered CA1 INs (see Methods). This analysis revealed that the first transcriptomic principal component (tPC1) primarily captures the global laminar position of INs, separating cells located in *stratum oriens* (green) from those in *stratum pyramidale* (magenta) (Fig. 4A, fig. S8C). Consistent with this, grouping CA1 INs by transcriptomic subclasses or types revealed a clear bifurcation along tPC1 based on their gross anatomical layer (fig. S8, D and F).

**Fig. 4.**
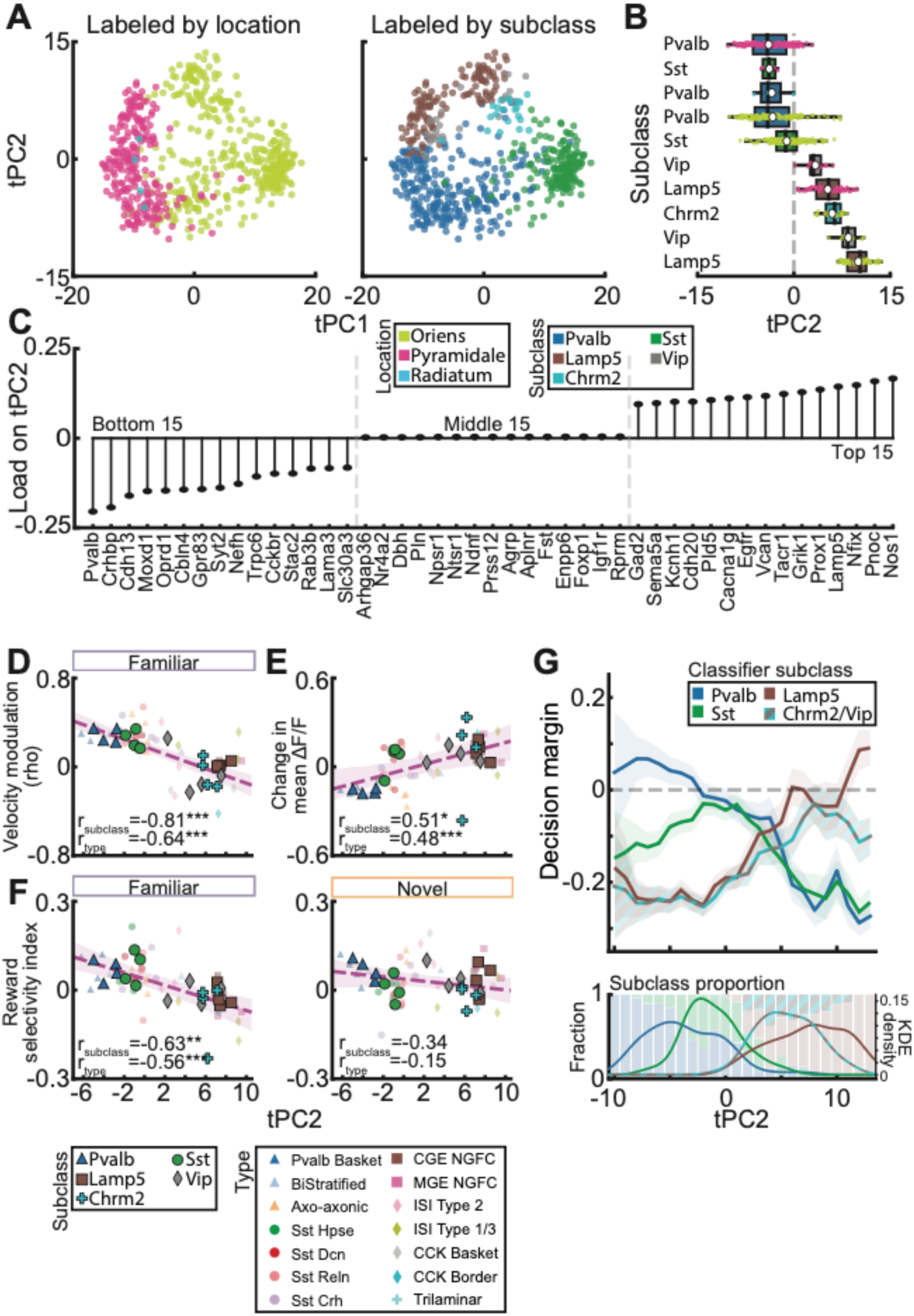
Organization of heterogeneous physiological responses along tPC2. (**A**) tPC2 versus tPC1 scores labeled by anatomical location (left) or transcriptomic subclass (right). tPC1 primarily reflects laminar position, whereas tPC2 captures the variation associated with subclass identity. (**B**) Distribution of CA1 IN subclasses along tPC2. Consistent with previous observations (*50*), transcriptomic subclasses occupy ordered positions along this axis. See *fig. S8* for type-level organization. Small colored circles indicate individual cells. Large white circles indicate the animal means. (**C**) Top, middle, and bottom 15 gene loadings contributing to the second transcriptomic principal component (tPC2). Gene loadings were obtained by running a dimensionality reduction on the cell-by-gene matrix of imaged neurons (see Methods). (**D**) Relationship between tPC2 and velocity modulation. tPC2 is strongly correlated with velocity modulation in the familiar context (shown here, r_subclass_=−0.81, p_subclass_=0.00018; r_type_=−0.64, p_type_=8.83×10^-7^) and in the novel context (see *fig. S9*). (**E**) Relationship between tPC2 and change in mean ΔF/F between contexts. tPC2 is significantly correlated with context-dependent changes in mean activity (r_subclass_=0.51, p_subclass_=0.032; r_type_=0.48, p_type_=0.0005). (**F**) Relationship between tPC2 and reward selectivity index in familiar (left) and novel (right) contexts. tPC2 score systematically varies with RSI in the familiar context (r_subclass_=−0.63, p_subclass_=0.0070; r_type_=−0.56, p_type_=0.00004), but this relationship is diminished in the novel context (r_subclass_=−0.34, p_subclass_=0.15; r_type_=−0.15, p_type_=0.32). (**G**) A random forest classifier trained exclusively on physiological features recapitulates the transcriptomic pattern of tPC2. Top: decision margin was defined in a one-vs-rest manner as the predicted probability of the indicated subclass minus the highest predicted probability among the remaining subclasses. For example, the *Pvalb* curve shows how much more likely a cell is to be classified as *Pvalb* than the most likely alternative subclass at each tPC2 position. *Chrm2* and *Vip* subclasses were merged because of low cell counts (n=43 cells). Positive margins indicate confident classification, whereas margins near zero indicate physiological ambiguity. The crossover of one-vs-rest decision margins along tPC2 provides independent support that subclass identity varies continuously with position along tPC2. Decision margins were binned into 1-unit tPC2 bins (24 bins), and the resulting curves were smoothed using a moving-window average (window=2 bins). Bottom: subclass fractions along tPC2 with kernel density estimates. For (**D**) through (**F**), smaller shaded symbols indicate type level, and larger bold symbols indicate subclass level. Shaded regions indicate 95% confidence intervals of the fitted line (purple), and p-values were corrected using the Benjamini-Hochberg FDR method for multiple comparisons. See table S4 for detailed statistics. For (**G**), shaded regions indicate the smoothed standard error of the mean (SEM).

In contrast, subclasses and types were distributed continuously along tPC2 (Fig. 4B, fig. S8, D and E). Consistent with previous reports for INs in the visual cortex (*50*), this axis did not simply recapitulate subclass boundaries but revealed a more complex structure. Notably, the AAC type clustered closer to the *Sst* subclass than to the *Pvalb* subclass, similar to observations in the visual cortex (fig. S8E) (*50*). Moreover, *Lamp5* and *Vip* types were intermingled at the opposite end of the spectrum with the MGE-derived NGFC type occupying the positive values along the continuum (Fig. 4B, fig. S8F). Consistent with this arrangement, tPC2 gene loadings spanned from *Pvalb-associated* genes to *Lamp5*-associated markers (Fig. 4C). Interestingly, tPC2 preserved fine-scale laminar positioning within individual subclasses and types (fig. S8D and E). Thus, while tPC1 captures the macro-scale anatomical locations of CA1 INs, tPC2 reveals a more continuous genetic gradient that aligns with subclass and type labels while retaining laminar information.

We next asked whether tPC2 was associated with the observed diversity in activity dynamics *in vivo*. To test this, we computed the mean tPC2 score and physiological features for each subclass within each animal, and calculated their correlation. For velocity modulation, we observed a significant negative correlation with tPC2, where subclasses with lower tPC2 scores exhibited stronger velocity modulation while those with higher tPC2 scores showed weaker modulation (Fig. 4D). This relationship was robust across contexts (Fig. 4D, fig. S9A), indicating that position along the tPC2 transcriptomic continuum tracked velocity modulation in both contexts.

We next examined whether tPC2 was related to context-dependent changes in mean ΔF/F activity (see Methods). The mean ΔF/F index had a positive correlation with tPC2, suggesting that CA1 INs at the lower end of tPC2 reduce their mean ΔF/F activity in the novel context (Fig. 4E). Surprisingly, while tPC2 tracked velocity modulation and mean ΔF/F signals, it did not significantly predict spatial tuning properties. Neither the rate map correlation nor change in sparsity (fig. S9, B and C) could be predicted by tPC2. Finally, we found that the relationship between reward selectivity index (RSI) and tPC2 was context-dependent. tPC2 was significantly correlated with RSI in the familiar context, but not in the novel context (Fig. 4F). These relationships were also observed at the type level (Fig. 4, D-F, fig. S9).

These results suggest that physiological properties are organized with respect to transcriptomic position along tPC2. To test whether this tPC2-associated structure was also reflected in the physiological feature space, we trained a random forest classifier to predict transcriptomically defined subclass labels using physiological features alone (fig. S10, A and B). Although the classifier was trained with discrete subclass labels, one-vs-rest decision margins shifted smoothly along tPC2, consistent with graded physiological similarity across subclass boundaries rather than abrupt transitions between subclasses (Fig. 4G, fig. S10, C and D). Together, these results suggest that a subset of physiological features is structured across both transcriptomically defined subclass and environmental context while physiology-based classifier margins provide orthogonal support for continuous ordering of subclasses along tPC2.

## Discussion

Here, we integrated *in vivo* two-photon Ca^2+^ imaging with *post hoc* MERSCOPE-based ST to establish a framework for directly linking detailed molecular identity with physiological profiles of inhibitory circuits in the mouse hippocampus (Fig. 1). This cross-modal pipeline enabled transcriptomic classification of GABAergic INs in CA1 into five major subclasses and 14 types, revealing laminar organization and relative abundances broadly consistent with prior scRNA-seq atlases (Fig. 2). During a VR context-switch task, these INs displayed heterogeneous, subclass-specific adaptations to novelty and reward (Fig. 3). This physiological diversity was correlated with position along a continuous transcriptomic axis (Fig. 4). This continuous organization extends findings from the neocortex, where transcriptomic gradients similarly predict electrophysiological properties and state modulation (*50, 51*).

This continuous transcriptomic axis preserved fine-scale within-type laminar organization in the hippocampus (Fig. 4B, fig. S8F), suggesting that laminar position and molecular composition provide complementary constraints on inhibitory circuit function in the hippocampus. Given the highly specialized input-output architecture of hippocampal laminae, the local excitatory environment in a given sublayer may further refine IN transcriptomic and functional properties. This is consistent with recent evidence in the neocortex showing that the composition and differentiation of *Pvalb* and *Sst* IN subtypes can be influenced by the proportion of their excitatory partners (*52*).

Although tPC2 correlated with certain physiological features such as velocity modulation and reward selectivity, it does not account for all physiological features of CA1 INs. Notably, it showed no significant correlation with spatial tuning properties, including rate-map correlations between contexts or changes in sparsity at the subclass level. Nevertheless, as in the neocortex (*50*), tPC2 may serve as an organizing principle for some of the functional heterogeneity observed among CA1 INs, while also representing an axis that orders CA1 IN subclasses and types. Furthermore, our dataset has limited representation of *Vip* and *Chrm2* subclasses, which constituted only ∼7% each of the registered imaged cells (n = 22 and 21, respectively), consistent with their low overall abundance in the full MERSCOPE dataset (∼14% and ∼3% of GABAergic INs, Fig. 2A). Given their position at the transition zone along tPC2, future studies employing subclass-specific Cre drivers (*53*) or AAV-enhancer (*54*) to enrich these populations will be valuable for testing the continuity and functional implications of the transcriptomic gradient.

Beyond the biological findings, our study establishes a scalable, robust multimodal pipeline that links gene expression to *in vivo* physiology at single-cell resolution. Recent studies have begun to relate transcript-based molecular identity to *in vivo* activity in cortical and subcortical circuits (*50, 55, 56*). To our knowledge, this is the first study to achieve multimodal, spatially resolved transcriptional-physiological mapping of hippocampal circuits during behavior. A key strength of this approach is that it preserves spatial information while enabling *post hoc* transcriptomic annotation of neurons recorded *in vivo*, allowing molecular identity, laminar position, and physiological function to be analyzed in the same hippocampal neuron. As spatial transcriptomic methods move toward whole-transcriptome coverage with improved spatial resolution (*4, 33, 57*), this framework may enable more comprehensive tests of how molecular programs organize hippocampal circuit function.

## Acknowledgments

We thank Drs. Franck Polleux, Ivo Spiegel, Boris Zemelman, and Allan-Hermann Pool for comments on an early version of the manuscript. Confocal imaging was performed with support from the Zuckerman Institute’s Cellular Imaging platform. We acknowledge the Vizgen technical support team for their assistance with troubleshooting the MERSCOPE protocols.

## Funding

Burroughs Wellcome Fund (SAH)

Boehringer Ingelheim Fonds PhD Fellowship (MCP)

National Institute of Mental Health grant R00MH128772 (EV)

National Institute of Mental Health grant R01MH124047 (AL)

National Institute of Mental Health grant R01MH124867 (AL)

National Institute on Aging grant RF1AG080818 (AL)

National Institute of Neurological Disorders and Stroke grant U01NS115530 (AL)

National Institute of Neurological Disorders and Stroke grant R01NS121106 (AL)

National Institute of Neurological Disorders and Stroke grant R01NS131728 (AL)

National Institute of Neurological Disorders and Stroke grant R01NS133381 (AL)

## Author contributions

Conceptualization: HCY, AL

Methodology: HCY, SAH, MCP, CKO, JY, BYR, TSM, JS, EV, AL

Investigation: HCY

Software: HCY, MCP, CKO, EV

Formal analysis: HCY, SAH, MCP, CKO, JY

Validation: HCY

Visualization: HCY, SAH, MCP, CKO, JY, SD

Data curation: HCY

Funding acquisition: AL

Resources: AL

Project administration: AL, EV

Supervision: AL, EV

Writing – original draft: HCY, SAH, MCP, CKO, JY, AL

Writing – review & editing: all authors

## Competing interests

Authors declare that they have no competing interests.

## Data and materials availability

Interactive example of the registration is available online at https://memorylongevity.org/merscope-registration. Data included in this study are available from the corresponding author upon reasonable request.

## Materials and Methods

### Subjects

All experiments were conducted in accordance with NIH guidelines and approved by the Columbia University Institutional Animal Care and Use Committee. Experiments were performed on four adult VGAT-Cre mice (Jackson Laboratory, RRID: 028862) to specifically label GABAergic interneurons in CA1. Sex was balanced in this study (two males and two females). Mice were housed with their littermates on a 12-hour light/dark cycle and had access to water *ad libitum* until the start of their behavior training.

### Surgeries

All surgeries were performed under anesthesia with isoflurane (3.5% for induction and 1.5% for maintenance) using a stereotaxic instrument (Kopf Instruments). Prior to incision, mice were given subcutaneous meloxicam and bupivacaine at the injection site. Body temperature was maintained with a heating pad during surgery and recovery. An eye ointment was applied to keep their eyes hydrated. Each animal received two surgeries, an injection and an implant, and these were separated by at least 3 days. Analgesics were provided as per Columbia University Institutional Animal Care and Use Committee prior to and after each surgery for up to 3 days.

For injections, mice aged 8-12 weeks were injected with 600 nL of a genetically encoded calcium indicator, AAV1-hSyn-FLEX-jGCaMP8s (Addgene: 162377-AAV1), using a glass pipette connected to a Nanoject III (Drummond Scientific). The vector was injected at six different locations to label GABAergic interneurons in different laminae of CA1 (AP: 2.25 mm, ML: 1.45 mm, DV: −1.55 to −1.05 mm; 100 nL per site, 100 µm steps). After the injection surgery, the incision site was closed using sutures.

At least three days after the injection surgery, a 1.7 mm craniotomy was made, centered around the injection site, and the cortex above CA1 was aspirated with continuous irrigation of cold 1× PBS. Aspiration was stopped when the medial-lateral fibers became visible to protect interneurons located in the most dorsal portion of CA1. A 1.7 mm cannula with a 1.7 mm glass coverslip was implanted to gain optical access to CA1 and secured in place with Vetbond. Finally, a custom titanium headpost was cemented to the skull using dental cement to head-fix the animals for two-photon functional imaging. Following the implant surgery, mice were allowed to recover for 7 days before beginning water deprivation for behavioral training.

### Behavior paradigm

Animals were trained to perform a spatial navigation task in a virtual-reality environment (*58*). Animals were first habituated to head fixation by placing them on the wheel for 15 minutes. The next day, animals were introduced to a 3 m track with 25 randomly positioned reward locations, which served as the familiar environment for the remainder of the experiment. Over subsequent training days, as animals learned to navigate the VR track, the number of rewards was progressively reduced to motivate running behavior. Once animals could complete 100 laps within 30 minutes with two fixed reward zones, they proceeded to imaging. During imaging, animals underwent a context-switch paradigm in which they were exposed in alternating blocks to the familiar environment and a novel environment that they had not previously encountered. Contexts were switched every 10 laps for a total of 60 laps. Reward zones were kept fixed in both environments. Context-switch experiments were conducted in two imaging sessions: one targeting *stratum oriens* and a second targeting *stratum pyramidale*. Different novel environments were used in the two sessions to ensure that animals were exposed to an environment they had never encountered before.

### *In vivo* two-photon imaging and calcium data preprocessing

Two-photon (2p) imaging was performed using an 8kHz resonant scanner (Bruker) and a 16x Nikon water immersion objective (0.8 NA, 3mm working distance). To image GCaMP8s signals, 820μm x 820μm field of view was imaged using a 920-nm laser (Coherent), and the power at the objective tip never exceeded 100mW. GCaMP signals were collected using a GaAsP photomultiplier tube detector (Hamamatsu, 7422P-40) following amplification via a custom dual-stage preamplifier (1.4 × 10⁵ dB, Bruker). All imaging was performed at 512 × 512 pixels with a digital zoom factor of 1. To image as many interneurons as possible, we performed multiplane imaging using a piezo objective scanner, with imaging planes spaced ∼30 μm apart. The frame rate was approximately 6 Hz per plane. As described above in the behavior paradigm section, context-switch experiments were conducted in two imaging sessions: one targeting stratum oriens and a second targeting stratum pyramidale. The two sessions were separated by at least 3 days.

Imaging data were first organized using the SIMA software package (*59*) and then processed with Suite2p (*60*) for motion correction and signal/neuropil extraction. ROIs were manually curated using Cellpose2 (*61*). After ROI curation, ROIs and their calcium transients were manually inspected in the Suite2p GUI. For ΔF/F computation, neuropil was subtracted from the raw fluorescence signal for each ROI. A static baseline was defined as the 7th percentile of the neuropil-corrected signal over the entire recording, and ΔF/F was computed relative to this baseline. The resulting ΔF/F traces were then smoothed using an exponential filter, and all analyses were performed on the smoothed ΔF/F traces.

### Regression of velocity-related activity

To control for the influence of velocity on interneuron activity, velocity was regressed out from neuropil-corrected signals using a ridge regression model. ΔF/F was then recalculated from the velocity-regressed traces using the same baseline and smoothing procedure described above. All analyses except those related to velocity modulation were performed using the velocity-regressed ΔF/F traces.

### Calculation of physiological features

Velocity modulation was quantified as the Pearson correlation between the animal’s running velocity and each interneuron’s ΔF/F. Prior to this correlation analysis, ΔF/F traces were max-normalized within each context.

For all analyses described below, frames with velocity < 5 cm/s were excluded. Mean ΔF/F in each context (familiar and novel) was computed by averaging ΔF/F across all frames within that context. For rate-map analyses, ΔF/F was binned into 5 cm spatial bins and smoothed with a Gaussian kernel (S.D. = 10 cm). Rate-map correlation was computed as the Pearson correlation between the average rate maps in the familiar and novel contexts. Split-half reliability was calculated by taking the average rate-map correlations between odd and even numbered trials within each familiar and novel context.

Sparsity was calculated as previously described (*62*). We computed the spatial sparsity of the rate map (with N spatial bins and r_n_ denoting the rate in the *n*th bin) as:

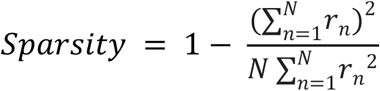

The reward selectivity index was computed as the difference between the mean ΔF/F activity in the two bins immediately preceding the first and second reward zone, normalized by their sum (positive values indicate preference for the first reward zone; negative values indicate preference for the second). The reward modulation index was computed as previously described (*13, 14*).

### Transcriptomics PCA

We performed generalized linear model principal component analysis (GLM-PCA; (*63, 64*)) on the cell-by-gene expression matrix from 610 imaged CA1 interneurons. All 500 genes in the panel were included in the analysis. Because GLM-PCA requires integer-valued inputs, we used the raw gene count matrix. We examined the resulting components and identified the second principal component (tPC2) as the axis that best captured the major continuum of transcriptomic variation across CA1 IN types. All downstream analyses used each cell’s scores along tPC2.

### Physiology-based classifier

To classify imaged interneurons based on physiological features alone, we trained a random forest classifier using 30 physiological features (table S5). Classification was performed at the subclass level. Because of the low number of cells in the *Vip* and *Chrm2* subclasses (n=43 cells in total), these subclasses were merged into a single class (*Vip/Chrm2*), resulting in four classes: *Pvalb*, *Sst*, *Lamp5*, and *Vip/Chrm2*. Each feature was percentile-normalized within each animal.

Classifier performance was evaluated using leave-one-animal-out cross-validation, in which the classifier was trained on three animals and tested on the held-out animal. To address class imbalance, we trained a bagged ensemble of 500 random forests using balanced downsampling. In each iteration, we sampled N_min_−1 cells per class without replacement, where N_min_ is the size of the smallest class. A random forest classifier was then trained using 600 trees, a maximum depth of 10, and a minimum leaf size of 5. Predicted class probabilities were averaged across iterations, and the final class label was assigned as the class with the highest mean predicted probability. Performance was summarized using accuracy and balanced accuracy.

The classifier margin shown in Fig. 4G was calculated in a one-versus-rest manner as the difference between the predicted probability of the indicated subclass and the highest predicted probability among the remaining subclasses. For example, the *Pvalb* curve indicates how much more likely a cell was to be classified as *Pvalb* than as its most likely alternative subclass at each position along tPC2.

### In vivo z-stack

On the last day of imaging, an *in vivo* z-stack was acquired to identify the same cells in both *in vivo* and *ex vivo* images. 4 fields of view were imaged with overlap to facilitate stitching, generating a large composite image that encompassed all areas that could be imaged with 2-photon imaging. To achieve a higher resolution than functional imaging, z-stacks were taken at 1024 × 1024 pixels with a digital zoom factor of 1.2 and a step size of 3-μm.

### Perfusion and tissue processing

A day after *in vivo* z-stack, intracardiac perfusion was performed with 20-25mL of cold 4% PFA. The brain was carefully removed from the skull, and fixed in 4% PFA overnight at 4°C. The next day, the brain was transferred to cold 15% sucrose, and kept for 8 hours. Then, it was transferred to 30% sucrose, where it remained until it sank, or for up to 36 hours. Finally, the brain was embedded in OCT, and frozen in prechilled 2-methylbutane (Millipore Sigma, M32631). It was then stored at −80°C until ready for processing in a cryostat.

### MERFISH experiment

All MERFISH experiments were performed using a fixed-frozen tissue processing protocol provided by Vizgen, with modifications. Briefly, brains were sectioned at 12-μm using a cryostat. After serial mounting, slides were air-dried in the cryostat for 1 hour and washed 3× in 1× PBS. To improve tissue adherence, slides were dried at room temperature for an additional 1 hour and then incubated overnight in 70% ethanol at 4°C.

The next day, *ex vivo* images were acquired before initiating MERSCOPE sample preparation. Sections were stained with DAPI (ACD RNAscope, 320858) and washed twice with Sample Prep Wash Buffer (Vizgen). All sections were imaged on a spinning-disk microscope (W1-Yokogawa inverted spinning disk confocal) using a 10x objective, acquiring both GFP and DAPI channels. Sample preparation was then resumed according to the Vizgen protocol. RNA pre-anchors from the FFPE kit were applied to improve RNA detection, as recommended by Vizgen. Sections were subsequently processed using the fixed-frozen protocol, including anti-rat and anti-chicken immunostaining to label GFP (rat monoclonal anti-GFP, Abcam, ab255886; chicken monoclonal anti-GFP, Abcam, ab300644) and cell boundary staining to aid segmentation. Following incubations and washes per the Vizgen protocol, sections were hybridized with the 500-gene panel for 48 hours. After washes per the Vizgen protocol, an acrylamide gel was applied and tissue was cleared for 14-18 hours to reduce background, following Vizgen recommendations. After clearing, samples were stained with DAPI and PolyT, then imaged on the MERSCOPE instrument with manually selected regions of interest.

### MERSCOPE image processing

MERSCOPE images were segmented using custom models trained with Cellpose 2 (*61*). The first model was trained using PolyT as the nuclear channel and cell boundary staining as the boundary channel, whereas the second model was trained using DAPI signal alone. For training, we generated small image snippets and manually segmented every cell within each snippet before training the models. Segmentation was performed at z-levels 3, 6, 9, and 12-μm using the Vizgen post-processing tool (VPT).

### Cross-modal registration across in vivo, ex vivo, and MERSCOPE images

The goal of the registration pipeline was to match individual cells across *in vivo* functional imaging, *ex vivo* section imaging, and MERSCOPE data within a common 3D coordinate framework. The final output consisted of CSV files linking MERSCOPE cell identifiers to the corresponding in vivo cell identifiers. An overview of the pipeline is provided in *fig. S1*. Registration was performed using a semi-automated, GUI-based framework with manual refinement, following a previously described approach (*44*). Briefly, registration was achieved through iterative selection of corresponding cells across imaging modalities, followed by sequential scaling, affine, and deformable transformations until satisfactory visual alignment was obtained.

To generate the *in vivo* reference volume, *in vivo* z-stack images acquired across four overlapping fields of view were stitched in x, y, and z using MATLAB custom code. This process included volume outline generation, image sharpening, spatial z-scoring to normalize brightness, padding, displacement-based estimation of relative tile placement (*65*), and final assembly into a stitched 3D volume. Both the stitched *in vivo* stack and each *ex vivo* section were then resampled according to imaging metadata so that voxel dimensions were isotropic in x, y, and z, downsampled isotropically, and saved for downstream registration.

Registration between *ex vivo* and *in vivo* datasets proceeded in two stages. First, each MERSCOPE section was aligned to its corresponding *ex vivo* section to account for differences in resolution, pixel size, and local tissue deformation, including non-isotropic shrinkage or expansion. For this step, the most in-focus MERSCOPE optical plane (z=4) was used. We downsampled MERSCOPE images in x and y (downsampling factor of 4) to accelerate computation. Second, the registered MERSCOPE and *ex vivo* images were combined into a two-channel image and each section was then independently registered to the stitched *in vivo* volume. Here, to perform registration, we first selected common cells across imaging modalities. Then, we applied scaling and affine transformation until satisfactory visual alignment was obtained. Furthermore, to this image, we continued to apply deformation and transformation iteratively, reducing Gaussian blur sizes. This step was repeated until their alignment was confirmed visually. The resulting transformations were applied to reconstruct 3D volumetric representations of the *ex vivo* and MERSCOPE datasets in the *in vivo* coordinate frame, thereby placing all three imaging modalities into a common reference space for subsequent mask matching. See *movie S1* for an example visualization of the registration process.

### Cross-modal cell matching

To align segmented masks across imaging modalities, imaging-session masks and *ex vivo* masks were separately mapped into the *in vivo* reference space. All segmented masks were generated using Cellpose2 or 3 (*61*) Because imaging-session masks were acquired as 2D images whereas the *in vivo* reference was a 3D stack, each imaging-session mask image was manually registered to the *in vivo* stack to identify the corresponding z-plane. The resulting transformations were then applied to the imaging-session masks, thereby placing them into the common in vivo coordinate frame.

*Ex vivo* masks were generated by first aligning each *ex vivo* section to its corresponding MERSCOPE section using DAPI signal. For this step, the most in-focus MERSCOPE optical plane (z=4) was used and downsampled in x and y to accelerate registration. The resulting transformation was applied to the *ex vivo* GFP image, after which a three-channel image consisting of registered *ex vivo* GFP, *ex vivo* DAPI, and MERSCOPE anti-rat signal was constructed for each section. These aligned multi-channel images were then manually segmented. The *ex vivo* masks were subsequently transformed through the previously computed registration steps so that they were represented in the same 3D coordinate frame as the *in vivo* stack. Malformed masks were excluded during processing.

To link *ex vivo* masks with MERSCOPE masks, *ex vivo* masks were mapped back into MERSCOPE coordinate space and compared with MERSCOPE segmentation masks. Candidate matches were identified based on spatial overlap (IoU > 0.25), and the best match was selected according to minimum centroid distance. This yielded *ex vivo* cell ID-MERSCOPE entity ID pairs. Finally, all modalities were visualized together in a Napari-based 3D interface, including the *in vivo* stack, transformed *ex vivo* volume, registered imaging masks, and registered *ex vivo* masks, to manually verify correspondence between *ex vivo* masks and imaging-session masks across imaging planes. These validated matches were combined with the *ex vivo*-MERSCOPE assignments to generate the final cell-matching tables linking *in vivo* functional masks to MERSCOPE masks. See *movie S1* for an example visualization of the registration process.

### Quality Control of MERFISH data

Cells were subjected to a comprehensive quality-control (QC) workflow (fig. S3, Ext. Data fig. 1A-G). Cells expressing fewer than 15 unique genes or fewer than 40 total transcripts were first discarded to remove poorly imaged cells. We then computed the fraction of each cell’s counts deriving from “Blank” control probes and excluded any cell with over 5% blank-probe contribution to ensure high-quality cells. Cells with volumes outside the 100–5,000 µm³ range were removed as well, and the remaining cells were normalized by volume (counts per µm), then globally rescaled to match the original mean transcript count for interpretability. To suppress the influence of extreme expression values, cells with total scaled counts falling outside the 1st–99th percentile were removed. Finally, doublets were detected using Scrublet (*66*) on the raw count matrix with up to 20 principal components. We applied a fixed doublet-score cutoff of 0.25 (consistent with previous MERFISH studies), and removed any cell exceeding this threshold. As we did not observe animal-dependent shifts in clustering during the analysis process, we did not require a harmonization step.

### Unbiased cell type clustering pipeline

Following QC, we performed hierarchical clustering using Scanpy (*67*). Counts were normalized per cell, log-transformed, and scaled. We computed principal components and constructed a k-nearest neighbor graph (k=10) using the top 20 PCs. To select optimal clustering resolutions, we performed stability analysis across 30–50 random initializations at resolutions ranging from 0.1 to 1.0, evaluating silhouette scores (cluster separation), number of clusters, and pairwise adjusted Rand index (ARI) across seeds (cluster stability). Cells were then clustered using the Leiden algorithm in three hierarchical levels: classes, subclasses, and types, with resolution parameters selected independently at each level to maximize silhouette score and ARI stability. UMAP embeddings were computed for visualization, and differentially expressed genes were identified using the Wilcoxon rank-sum test (adjusted p < 0.05).

### Assignment of subtype identities based on MERSCOPE transcriptomic profiles

Interneuron subtype names were assigned based on our unbiased cell type clustering (Extended Data figure 1) and validation pipeline described above and final manual annotation in reference to published cell type-specific studies for hippocampal interneuron cell types and single-cell sequencing based atlases atlases (*1, 10, 36, 45, 68–70*). From our iterative reclustering, we defined fourteen molecularly distinct IN types identified by differential combinatorial marker gene expression. Among MGE-derived populations, the *Pvalb* subclass was segregated into basket, axo-axonic, and bistratified cells (Ext. Data fig. 1, O-S). The *Sst* subclass was split into four types distinguished by differential expression of *Reln*, *Crh*, *Hpse*, *Dcn*, and *Npas1* (Ext. Data fig. 1, T-X). The *Lamp5* subclass comprised both CGE- and MGE-derived NGFCs (*71*) (Ext. Data fig. 1, Y-CC). Within the CGE-derived *Vip* subclass, we identified cell clusters consistent with *Cck*-expressing basket and border cells as well as two IN-selective IN subtypes (Ext. Data fig. 1, DD-HH). The *Chrm2* subclass did not cluster further and consists of the single trilaminar cell type. Below we detail our complete resulting taxonomy and top differentially expressed genes for each clustering level (class, subclass, and cell type):

***Class level taxonomy****: All cells, top differentially expressed genes (DEGs)*

**CA1 Pyramidal Neurons:** *Wfs1, Slc17a7, Chrm1, Grin2a, Elmod1*
**Astrocytes:** *Sox9, Aldoc, Ntsr2, Gfap, Daam2*
**Oligodendrocytes & Oligodendrocyte Progenitor cells (OPCs):** *Olig1, Sulf2, Ddr1, Olig2, Ndrg1*
**Vascular Cells:** *Cldn5, Flt1, Igf1r, Ptprm, Rgs12*
**GABAergic Interneurons:** *Slc32a1, Gad1, Gad2, Sp9, Rab3b*
***Subclass level taxonomy:*** *GABAergic Interneurons, top DEGs*

**Pvalb:** *Pvalb, Nefh, Oprd1, Nefm, Kcnc3*
***Sst:*** *Sst, Crhbp, Grin3a, Elfn1, Barx2*
***Lamp5:*** *Nfix, Nos1, Gad1, Gad2, Lamp5*
***Vip:*** *Npas1, Cnr1, Fxyd6, Trp53i11, Cck*
***Chrm2:*** *Chrm2, Cpne4, Grik1, Alk, Ptpru*
***Cluster level taxonomy:*** *GABAergic interneuron cell types in our study and justification*

#### Pvalb subclasses

##### Pvalb Basket

Parvalbumin-expressing basket cells are abundant pyramidal neuron soma-targeting inhibitory interneurons that primarily reside in the *stratum pyramidale*. (*72–74*) They are identifiable by their relative enrichment of *Pvalb* as well as *Syt2* expression (*75, 76*). Top DEGs: *Syt2, Nefh, Gad1, Kcnc3, Pvalb, Gpr83*.

##### Axo-axonic

Also known as Chandelier cells due to their finger-like axon projections into the pyramidal layer, Axo-axonic cells exhibit specific contact for the axon initial segment of CA1 pyramidal cells (*38, 73, 74*). Among *Pvalb*-expressing cells they are unique in this subclass for *Unc5b* expression, which has been validated as a reporter gene for these cells (*77*). Top DEGs: *Pthlh*, *Unc5b, Adgra1, Kcnip2, Chrm1*.

##### BiStratified

BiStratified cells are bipolar dendrite targeting cells of the hippocampus and neocortex (*73, 74*). These cells are identified by their increased expression of *Sst* (*78*) and *Npy* among high *Pvalb*-expressing cells (Ext. Data fig. 1, Q-S). (*79*). Top DEGs*: Sst, Npas1, Vgf, Npy*.

#### Sst subclasses

Oriens-Lacunosum Moleculare (O-LM) targeting *Sst*-expressing cells are a dominant cell type amongst the *stratum oriens*. These abundant dendrite targeting cells are typically identified morphologically by their soma localization in the *stratum oriens* of CA1 and their unique targeting of distal CA1 pyramidal cells dendrites in the *stratum lacunosum moleculare* layer (*36, 78, 80, 81*). However, transcriptomic markers to further subdivide this subclass have not been well investigated, and the 500-gene PanNeuro panel from Vizgen (table S6) did not include several key genes that could have enabled their characterization based on recent studies (*82*), nor do we have cell morphology data that might have assisted in our classification. Thus, we clustered our *Sst*-expressing subclasses using our clustering integrity parameters, yielding four *Sst* clusters, which we named after the top individual genes differentially expressed between clusters from our gene panel (Ext. Data fig. 1, T-X).

**Sst Crh**

Top DEGs: *Sst, Crh*

**Sst Dcn**

Top DEGs: *Sst, Dcn, Npy*

**Sst Hpse**

Top DEGs: Sst, *Hspe, Crhbp, Pou4f1*

**Sst Reln**

Top DEGs: *Sst, Reln*

#### Lamp5 subclasses

##### MGE NGFC

Medial ganglionic eminence (MGE)-derived Neurogliaform (NGFCs) and Ivy Cells. Ivy cells are abundant MGE-derived INs that fall into the *Lamp5* subclass of dendrite-targeting cells, morphologically identifiable by their English ivy-like appearance of their axons that densely branch close to their origin (*71, 83*). They have only been described in the hippocampus, densely targeting proximal basal and oblique dendrites of pyramidal cells in the *stratum oriens* and *stratum pyramidale* (*83–85*). They are transcriptomically differentiated from CGE NGFCs by differential expression of *Lhx6*, *Nos1*, and completely unique expression of *Tacr1*(*1, 83*). Top DEGs: *Tacr1*, *Ptprt, Rasgrf2, Nos1, Lhx6*.

##### CGE NGFC

Caudal ganglionic eminence (CGE)-derived neurogliaform cells. Neurogliaform cells are both MGE and CGE-derived INs that fall into the *Lamp5* subclass of dendrite-targeting cells. Their somas reside in the *stratum radiatum* and *stratum lacunosum moleculare* densely targeting distal oblique dendrites of CA1 pyramidal cells in those same regions (*86–89*). These cells are histologically determined by their -nNos expression along with other positive markers for neurogliaform cells with immunohistochemistry in tissues, consistent with decreased transcriptomic expression of *Nos1* in this *Lamp5* cluster (*71*). Top DEGs: *Chst1*.

#### Vip subclasses

##### ISI Type 1/3

Interneuron-Selective Interneurons Type 1 & Type 3 cells. This cluster represents CGE-derived disinhibitory GABAergic neurons expressing high *Vip* and *Calb2,* primarily targeting GABAergic neurons including *Sst*-expressing O-LMs and Cck Basket cells, and less commonly targeting BiStratified and Pvalb Basket cells (*36, 87–89*). Typically types 1 and 2 are separable by ISI Type 1 absence of *Vip* expression immunohistochemically (*90*); however, this cluster did not resolve these two types, either by differences between transcriptomic and proteomic detection or total number of cells collected, and thus likely it is a combined grouping of both. Top DEGs: *Calb2*, *Grin3a, Parm1, Rgs12, Npy2r*.

##### ISI Type 2

Interneuron-Selective Interneurons Type 2 cells. These CGE-derived cells target GABAergic dendrites restricted to stratum lacunosum-moleculare. This cluster was identified by its high expression of *Vip* and its absence of *Calb2* and *Cck* expression (*36, 87, 89*). Top DEGs: *Vip, Grin2a, Nefm, Elfn1*.

##### Cck Basket

Cholecystokinin-expressing basket cells. These are CGE-derived, pyramidal cell soma-targeting basket cells historically identified by their double labeling of Cck and Vip protein expression (*87, 89*) as well as a cannabinoid receptor (*Cnr1*) (*91*). More recently through transcriptomic atlases *Sncg* has been identified as a unique marker for targeting these cells (*1*). They exhibit an alternating perisomatic inhibition with Pvalb Basket Cells, which inhibit Cck Basket cells directly (*91, 92*). Top DEGs: *Sncg, Trp53i11, Cnr1, Sp8, Cck*.

##### Cck Border

Cholecystokinin border cells. Cck Border cells are CGE-derived dendrite targeting Cck-expressing interneurons that largely reside in *stratum radiatum* and enrich at the border with *stratum lacunosum moleculare* (*93–95*). Interestingly, we found that *Col14a1* expression is unique to this cluster, which mirrors single cell atlas data (*1*). Top DEGs: *Col14a1, Ptprm, Sp8, Cnr1, Sulf2*.

#### Chrm2 *subclass and* Trilaminar *type*

We identified these cells from the GABAergic interneuron subclass level as subsequent clustering resulted in a homogenous population that did not further cluster (Extended Data fig. 1, L-N). Given their high muscarinic receptor expression (*Chrm2, Chrm4*) combined with *Sst* expression, along with their spatial locations restricted to the border of the *alveus* and *oriens*, these cells may include the long-projecting theta-off ripple-on (TORO) cells of the hippocampus and Trilaminar cells (*53, 96, 97*). Top DEGs: *Chrm2, Cpne4, Grik1, Alk, Ptpru*.

### MapMyCells reference annotation to Allen Brain Cell Atlas

Cell Type Mapper (MapMyCells; RRID:SCR_024672) was used to map our cells to the Allen Brain Cell Atlas taxonomy. MERFISH dataset of a single adult mouse brain (MERFISH-C57BL6J-638850 (*2*) from Allen Brain Cell Atlas was used to build a reference tree. Hippocampal cells from the atlas were selected by ‘*cell_metadata_with_parcellation_annotation’* from MERFISH-C57BL6J-638850-CCF (parcellation_division= ‘HPF’). WMB-taxonomy metadata was used to link ‘*cluster membership’* and ‘*cluster_alias’*. After subsetting the dataset to hippocampal region/cells, three prerequisite .h5 files were generated (1) The precomputed stats file (2) reference markers (3) the query markers. 188 out of 500 genes from our MERFISH gene panel overlapped with the reference atlas.

### Statistical analysis

All statistical details are provided in the figure captions. No statistical methods were used to predetermine sample sizes. Boxplots show the 25th, 50th (median), and 75th percentiles, with whiskers extending to 1.5 times the interquartile range below the 25th percentile or above the 75th percentile. Outliers are defined as values beyond the whisker range. Unless otherwise specified, paired Student’s t-tests were performed for each subclass. Type-level analyses were performed only when the corresponding subclass-level t-test was significant. For type-level analyses, p-values were corrected for multiple comparisons using the Benjamini–Hochberg false discovery rate procedure within each type.

---

**Fig. S1.**
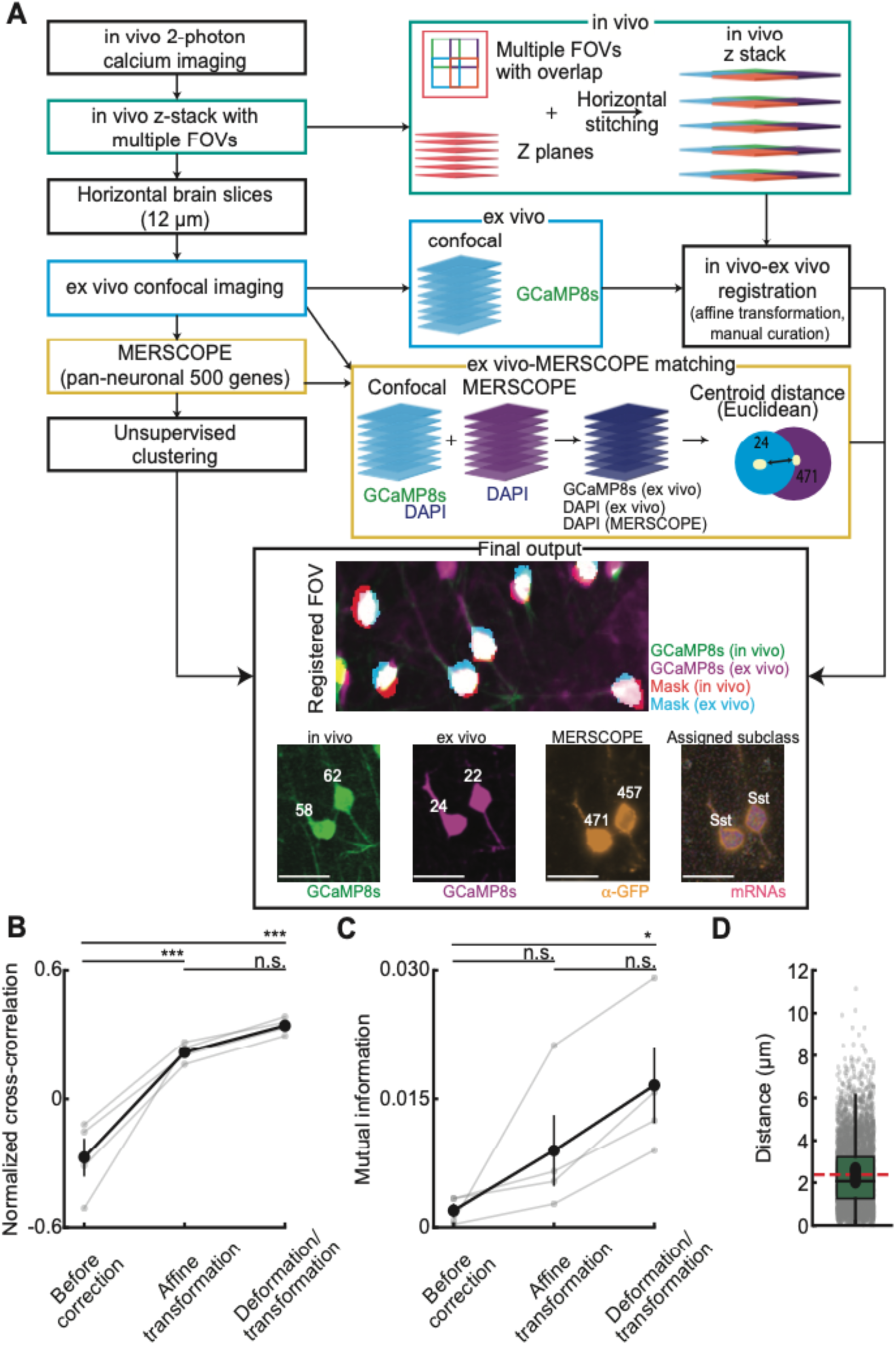
Experimental pipeline and registration metrics. (**A**) Schematic of the full experimental pipeline. After the *in vivo* two-photon calcium imaging, high-resolution z-stacks were acquired. Multiple z-stacks were acquired with overlapping fields of view (FOVs) and stitched to generate a large composite image that matches the *ex vivo* tissue area. Brains were horizontally sectioned (12-µm thickness) and imaged using confocal microscopy, followed by MERSCOPE processing. *Ex vivo* images were registered to *in vivo* z-stacks using a sequence of deformations and transformations, and manual curation. To align MERSCOPE data, *ex vivo* GCaMP fluorescence was segmented, and MERSCOPE DAPI images were registered to *ex vivo* DAPI. The same cells were identified by first selecting candidates with segmentation overlap (intersection over union (IoU)>0.25), and assigning matches based on minimal centroid distance. As a final verification step, segmented *ex vivo* GCaMP+ cells and *in vivo* cells were manually confirmed as matches in their registered 3D space. Transcriptomic cell types were assigned via unsupervised clustering of the MERSCOPE spatial transcriptomics data. See Methods for details and refer to *movie S1* for an additional visualization of the registration process. (**B**) Normalized cross-correlation between *in vivo* and *ex vivo* images significantly improved across registration steps (one-way repeated measures ANOVA, F(2,6)=36.24, p=0.0004). Post-hoc analysis confirmed that the final deformation/transformation step yielded a significant improvement over the uncorrected state (p=0.001, Tukey HSD). (**C**) Mutual information between *in vivo* and *ex vivo* images showed a significant improvement across registration steps (F(2,6)=7.48, p=0.024). *Post hoc* tests confirmed that the final deformation/transformation step yielded a statistically significant improvement over the uncorrected state (p=0.04, Tukey HSD). (**D**) Euclidean distances between the centroids of matched *ex vivo* and MERSCOPE segmented masks. Individual cells from all animals are shown in gray overlaid on the boxplot. Black circles indicate animal means, and the red line indicates the grand mean across animals. For (**B**) and (**C**), gray lines indicate individual animals, while the black line indicates the animal mean ± SEM. See *table S1* for detailed statistics.

**Fig. S2.**
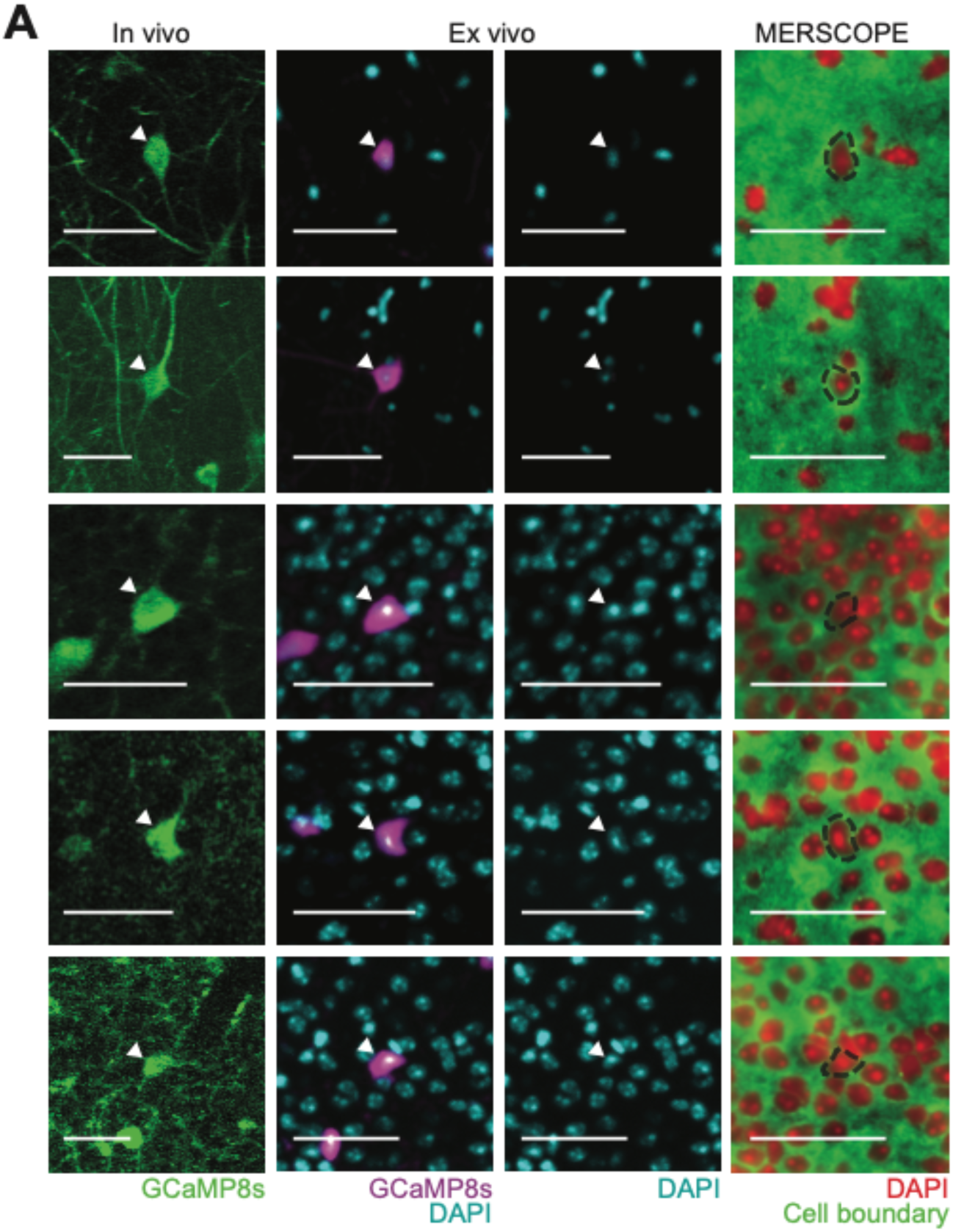
Cross-modal registration of imaged CA1 INs. Examples of a single CA1 IN (arrowheads) tracked from *in vivo* calcium imaging (left column) to *ex vivo* imaging (middle columns; GCaMP8s in magenta, DAPI staining in cyan) and MERSCOPE (right column; DAPI in red, cell boundary staining in green). Black dashed outlines indicate MERSCOPE segmentation masks of matched nuclei (see **Methods**). The scale bar is 50-µm. Refer to *movie S1* for an additional visualization of the registration process.

**Fig. S3.**
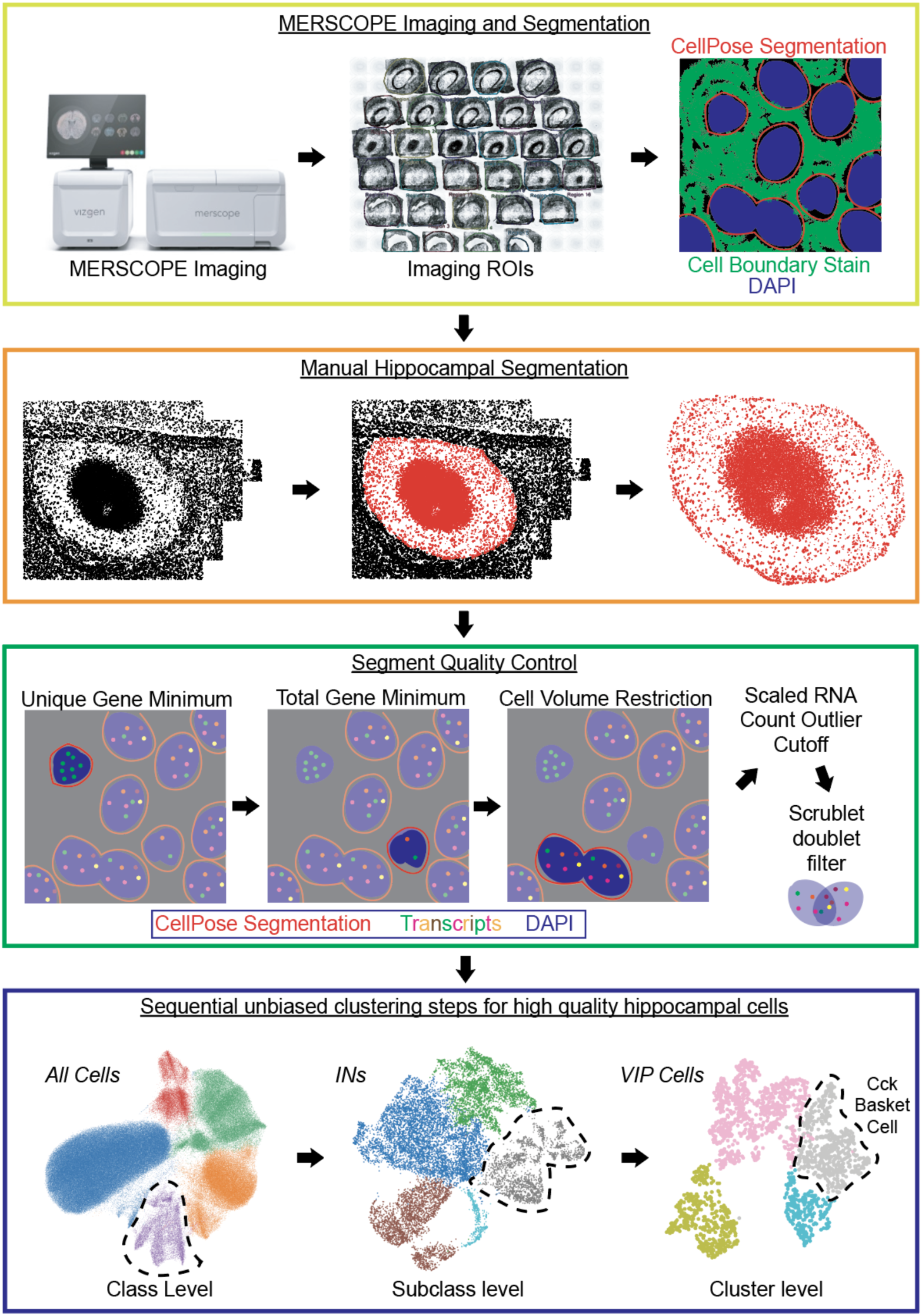
Overview schematic of unsupervised cell classification scheme for GABAergic INs. ∼25 x 12 μm transverse sections are taken from the *in vivo* imaging area under the cannula and carefully aligned within the MERSCOPE imaging window on a single cover slip. The MERSCOPE imaging and analysis platform from Vizgen performs both imaging and segmentation analysis, accounting for cell boundary stains and the DAPI signal for cell segmentation with Cellpose (top). Segments outside hippocampal regions in the output, visualized in the Vizgen Visualizer software, are manually removed from the dataset (middle top). Several quality-control steps are implemented to remove low-quality cells, duplicates, and nonsense segments from the dataset (middle and bottom). Finally, sequential unbiased clustering steps are performed with quality-control measurements at three levels: class level for major cell-type categories, subclass level for major subtypes of GABAergic IN classification, and cluster level for individual GABAergic interneuron cell types (bottom). Cell types are assigned canonical naming conventions following the literature. *ROIs*: Region of interest. *INs*: GABAergic interneurons. *VIP cells*: Vasoactive intestinal peptide-expressing cells. *Cck*: Cholecystokinin-expressing cells.

**Fig. S4.**
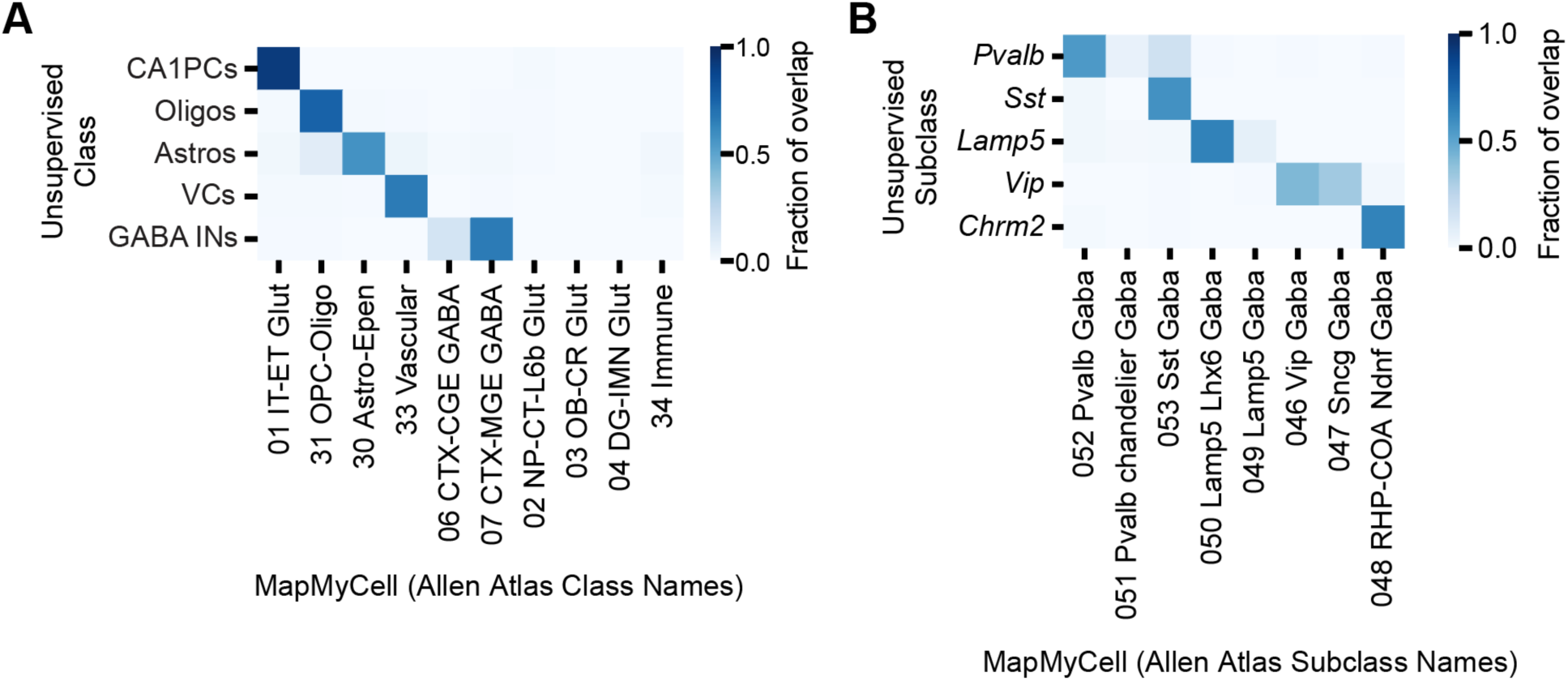
Jaccard similarity of unsupervised clustering with Allen Brain Cell Atlas data. (**A**) Fraction of cells per unsupervised clusters matched to Allen Brain Cell Atlas taxonomic classes. (**B**) Fraction of cells per subcluster within GABA INs class matched to Allen Brain Cell Atlas subclasses. CA1PCs: CA1 pyramidal cells. Oligos: Oligodendrocytes and oligodendrocyte progenitors. Astros: Astrocytes. VCs: Vascular cells. GABA INs: GABAergic inhibitory interneurons.

**Fig. S5.**
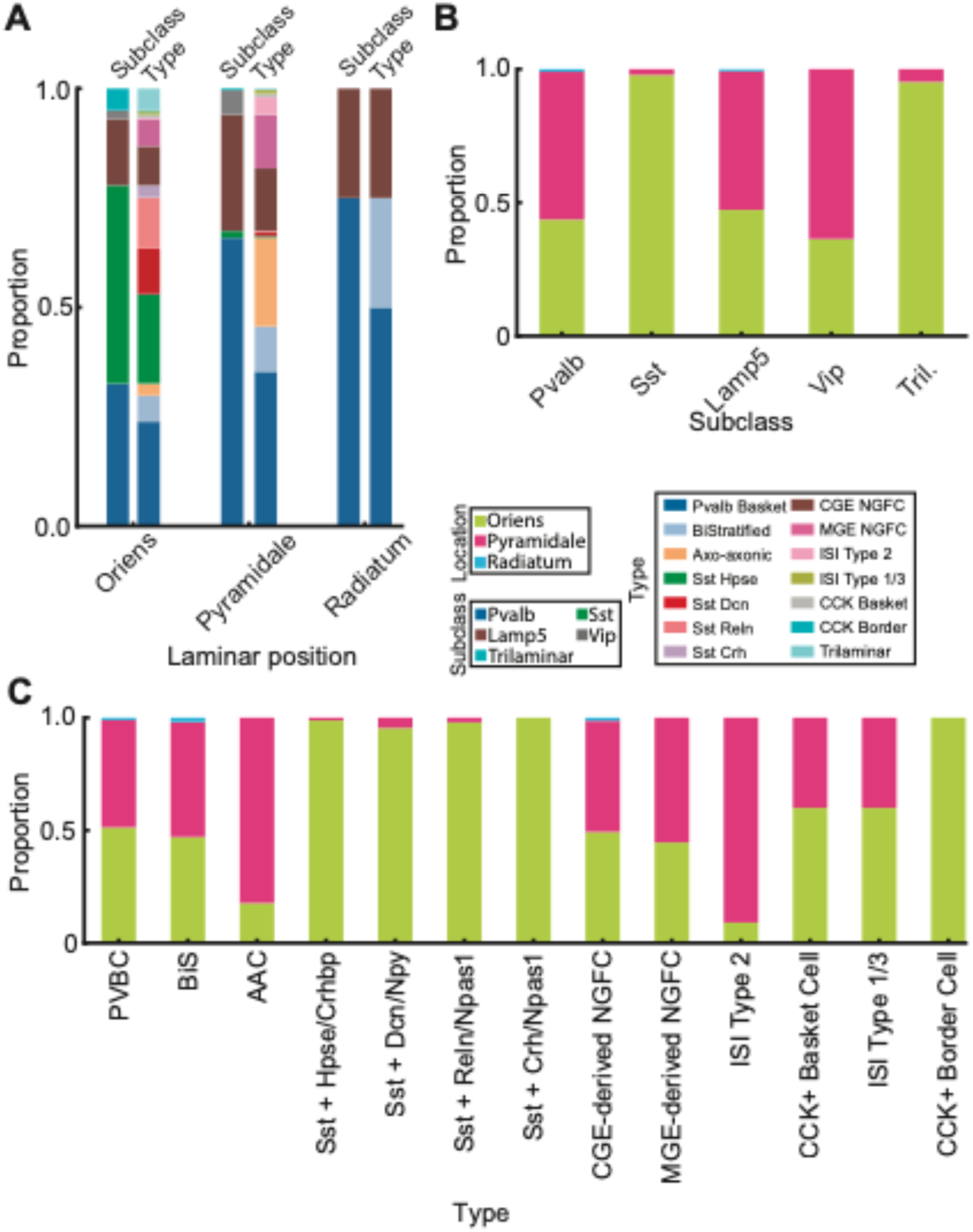
Laminar organization of registered CA1 INs. (**A**) Proportion of registered CA1 INs identified within the *stratum oriens*, *pyramidale*, and *radiatum*, classified by transcriptomic subclass (left) and type (right). Colors correspond to the subclass and type legends. See *tables S2 and S3* for quantification. (**B**) Laminar distribution of the five major IN subclasses. Stacked bars show the proportion of each subclass in the *stratum oriens* (green), *pyramidale* (magenta), and *radiatum* (blue). (**C**) Laminar distribution of transcriptomic types. Data represent all imaged and registered INs (n=610 from 4 mice).

**Fig. S6.**
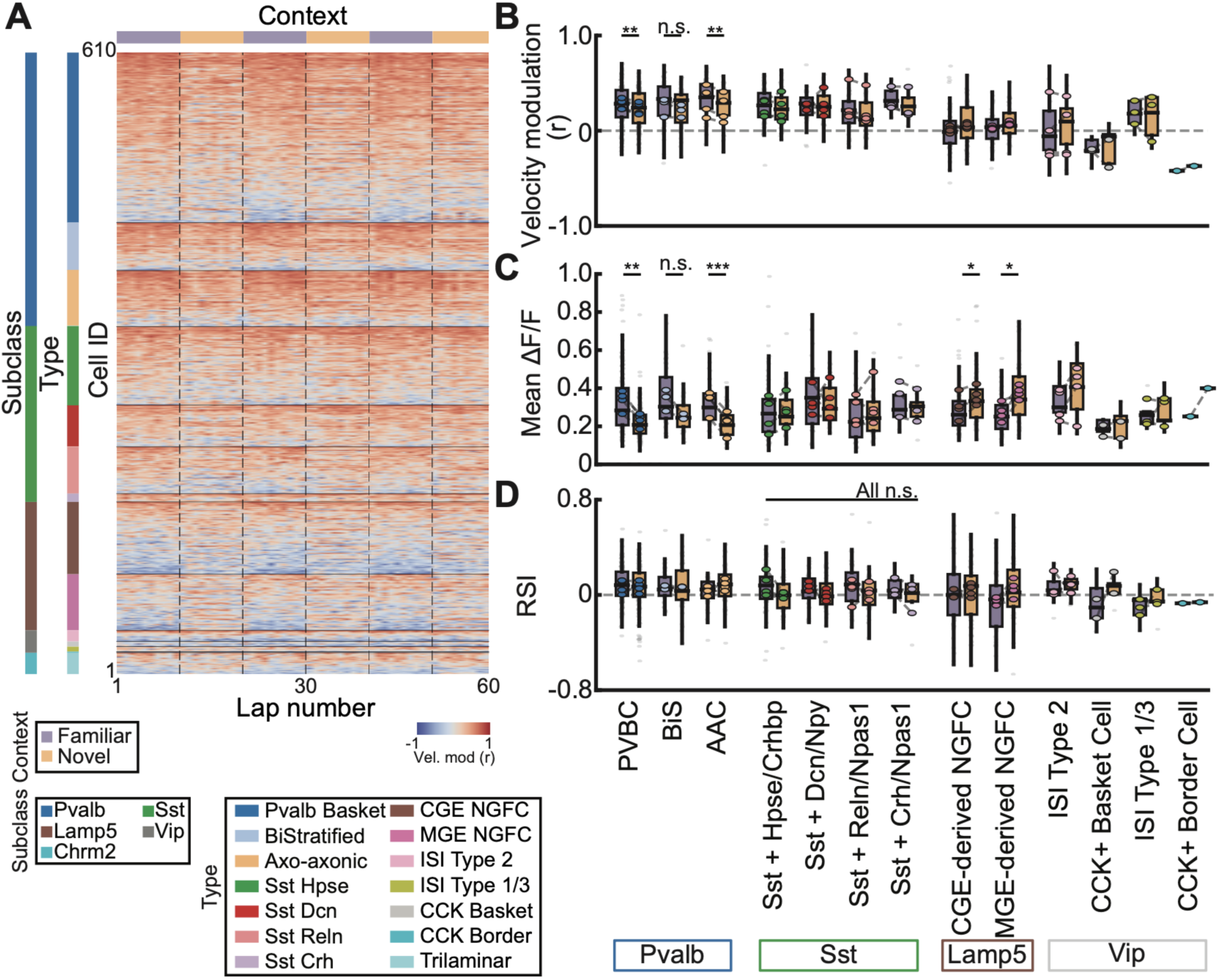
Type-specific physiological features across contexts. (**A**) Lap-by-lap velocity modulation for individual CA1 INs during the context switch task. Rows represent individual CA1 INs grouped by their subclass. They were further sorted within type by their mean velocity modulation during the first block of the familiar context. (**B**) Velocity modulation of CA1 INs by transcriptomic types in familiar (purple) and novel (orange) contexts. *Pvalb* Basket cells (PVBC) (t(3)=6.91, p=0.0093, n=4 animals) and AAC (t(3)=8.74, p=0.0032, n=4 animals) showed significant context-dependent changes. *Pvalb* BiStratified (BiS) was not significantly different (t(2)=2.04, p=0.18, n=3 animals). (**C**) Changes in mean ΔF/F across contexts. PVBC (t(3)=8.60, p=0.0050, n=4 animals) and AAC (t(3)=21.62, p=0.00022, n=4 animals) showed a significant reduction in mean ΔF/F in novel contexts while BiS did not (t(2)=3.09, p=0.091, n=3 animals). Both CGE-NGFC (t(3)=−3.97, p=0.029, n=4 animals) and MGE-NGFC (t(3)=−4.47, p=0.029, n=4 animals) types significantly increased their mean ΔF/F in the novel context. (**D**) Reward selectivity index (RSI) across contexts. Sst types did not show significant changes in RSI between contexts (Sst-Hpse: t(3)=2.33, p=0.27, n=4 animals; Sst-Dcn: t(3)=2.02, p=0.27, n=4 animals; Sst-Reln: t(3)=0.053, p=0.96, n=4 animals; Sst-Crh: t(2)=1.45, p=0.38, n=3 animals). P-values from paired t-tests were corrected with Benjamini-Hochberg FDR. Statistical quantification was restricted to types that had significant changes at the subclass level. For panels (**B**) to (**D)**, boxplots indicate the distribution of individual cells (gray). Colored circles represent animal means.

**Fig. S7.**
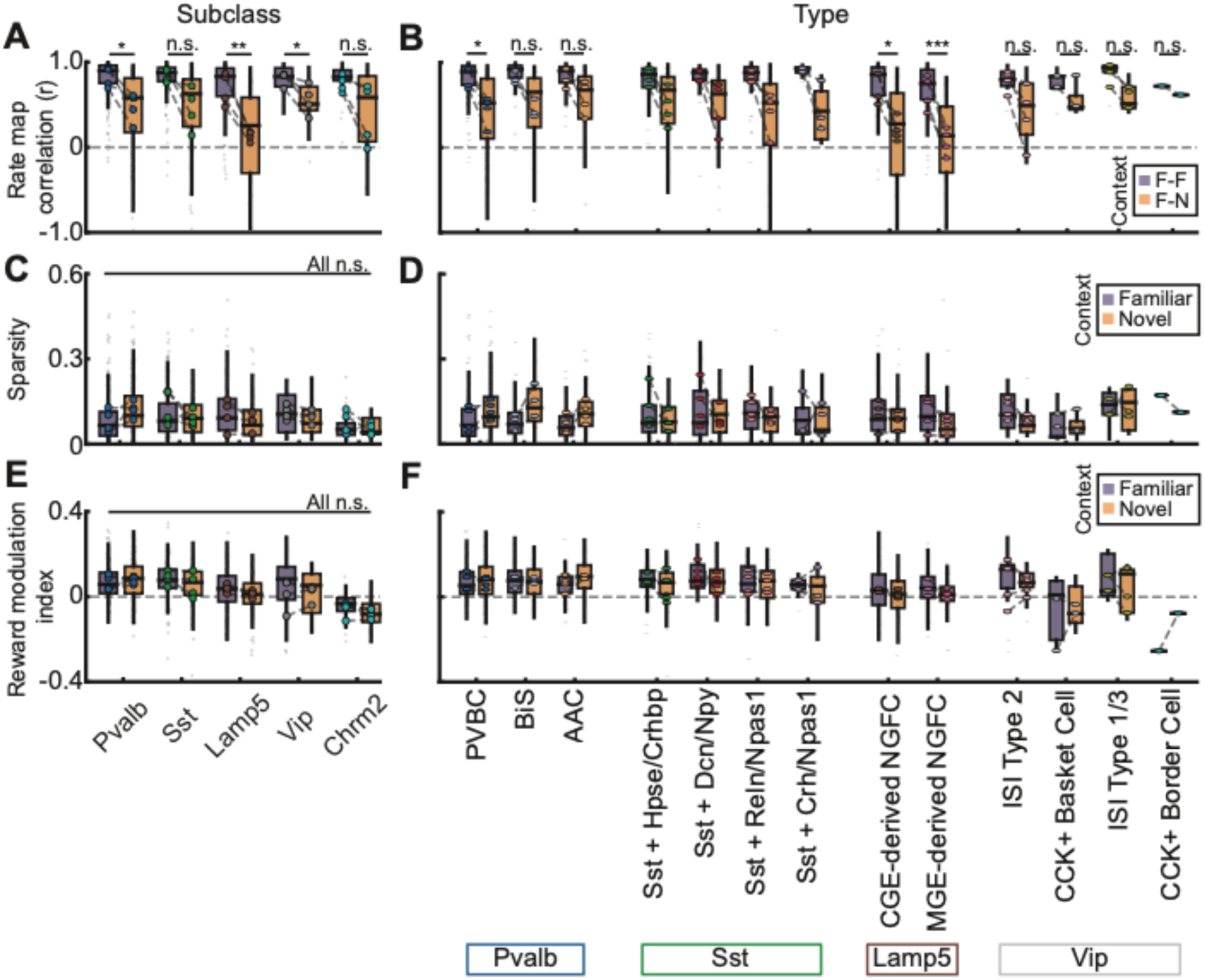
Additional physiological features of CA1 INs across contexts. (**A**) Rate map correlation across contexts at the subclass level. Rate map correlation between familiar and novel contexts (F-N) was compared with familiar split-half correlation (F-F). Significant differences were observed for *Pvalb* (t(3)=4.90, p=0.016, n=4 animals), *Lamp5* (t(3)=8.04, p=0.0040, n=4 animals), and *Vip* (t(3)=4.27, p=0.024, n=4 animals). (**B**) Same analysis at the type level. F-N rate map correlation differed significantly for PVBC (t(3)=5.42, p=0.037, n=4 animals), but not for BiS (t(2)=3.84, p=0.066, n=3 animals) or AAC (t(3)=2.83, p=0.066, n=4 animals). Within the *Lamp5* subclass, both MGE-NGFC (t(3)=9.33, p=0.0026, n=4 animals) and CGE-NGFC (t(3)=4.20, p=0.025, n=4 animals) showed significant differences. Within the *Vip* subclass, no types remained significant after FDR correction. (**C**) Sparsity at the subclass level. No subclasses showed significant changes in sparsity between contexts. (**D**) Sparsity at the type level. (**E**) Reward modulation index (RMI) at the subclass level. No subclasses showed significant changes in RMI between contexts. (**F**) Same as (E), but at the type level. P-values from paired t-tests were corrected with Benjamini-Hochberg FDR. Statistical quantification was restricted to types that had significant changes at the subclass level. For all panels, boxplots indicate the distribution of individual cells (gray). Colored circles represent animal means.

**Fig. S8.**
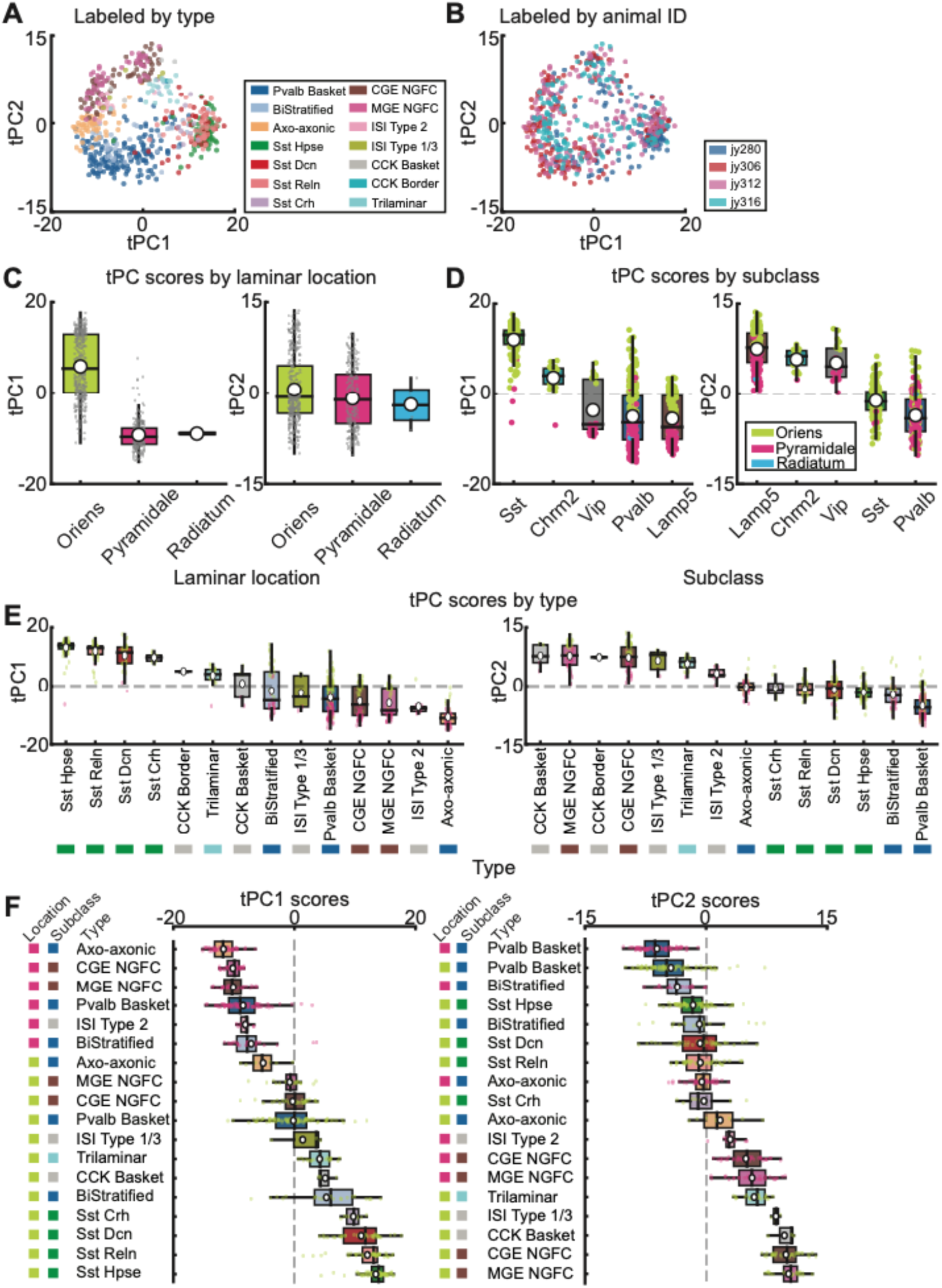
Characterization of transcriptomic principal components (tPCs). (**A**) Distribution of registered CA1 INs in tPC space (tPC1 vs. tPC2) colored by type. (**B**) Same as panel (**A**), but colored by animal ID. Note the lack of batch effects. **(C**) tPC1 (left) and tPC2 (right) scores separated by laminar location of registered CA1 INs (n=610 cells). tPC1 primarily captures the laminar position, separating *stratum oriens* from *pyramidale*. (**D**) tPC1 (left) and tPC2 (right) scores of registered CA1 INs separated by subclass labels (n=610 cells). (**E**) Same as panel (**D**), but separated by type labels. Groups are ranked by animal means. Note the progression of types along the tPC2 axis while tPC1 largely separates the laminar location. (**F**) tPC1(left) and tPC2 (right) scores separated by type and laminar location. For all panels, scatters indicate individual cells. For panels (**C**) to (**F**), boxplots indicate the distribution of individual cells. White circles represent animal means.

**Fig. S9.**
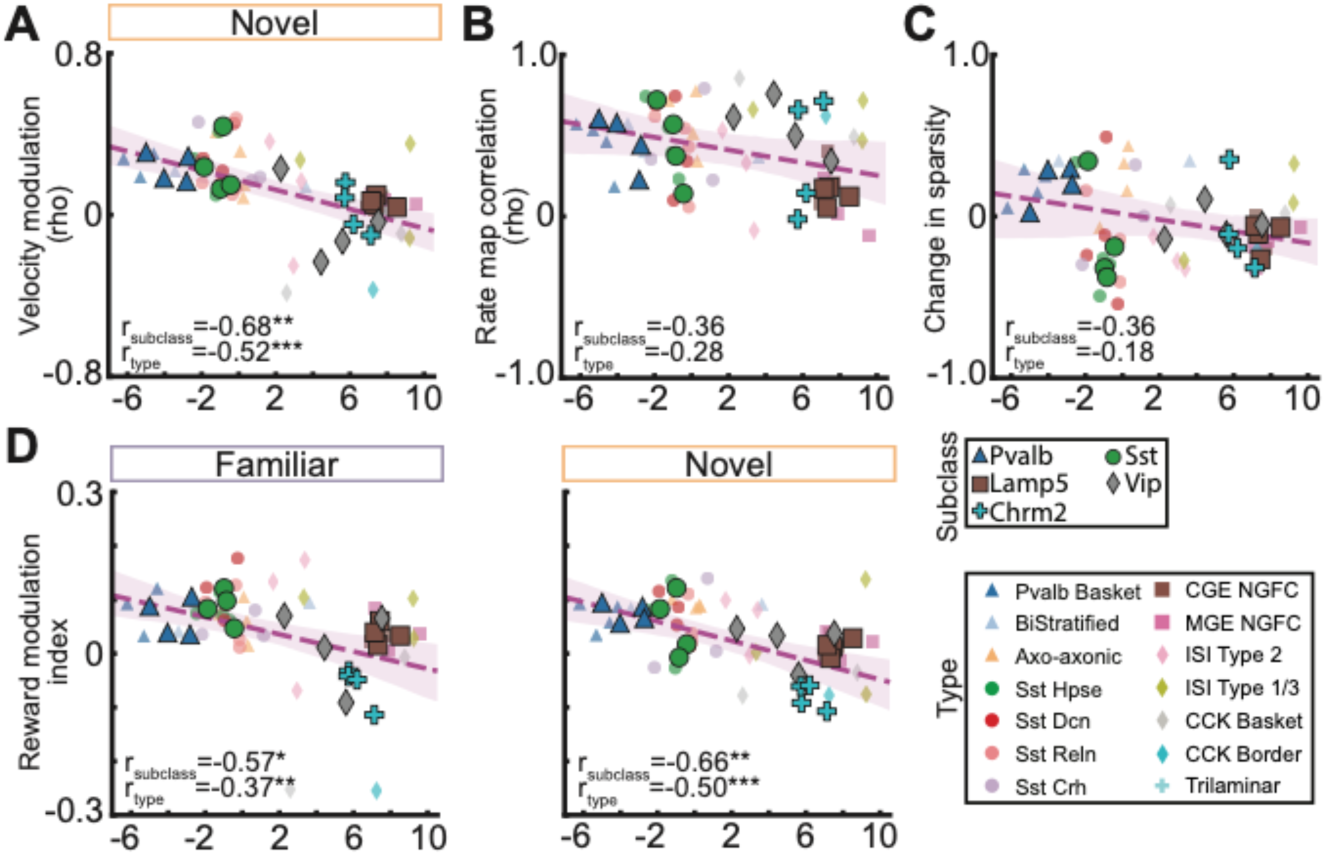
Correlation between tPC2 and physiological features. (**A**) Correlation between velocity modulation and tPC2 scores (r_subclass_=−0.68, p=0.0046; r_type_=−0.52, p_type_=0.00015, FDR-corrected). (**B**) Correlation between rate map correlation across contexts and tPC2 scores (r_subclass_=−0.36, p_subclass_=0.13; r_type_=−0.28, p_type_=0.059, FDR-corrected). (**C**) Correlation between change in sparsity (r_subclass_=−0.36, p_subclass_=0.13; r_type_=−0.18, p_type_=0.21, FDR-corrected). (**D**) Reward modulation index in the familiar (r_subclass_=−0.57, p_subclass_=0.016; r_type_=−0.37, p_type_=0.0094, FDR-corrected) and novel (r_subclass_=−0.66, p_subclass_=0.0049; r_type_=−0.50, p_type_=0.00032, FDR-corrected) contexts. Smaller shaded symbols indicate type level, and larger bold dots indicate subclass level. Shaded regions indicate 95% confidence intervals of the fitted line (purple). *p < 0.05, ** p < 0.01, *** p < 0.001

**Fig. S10.**
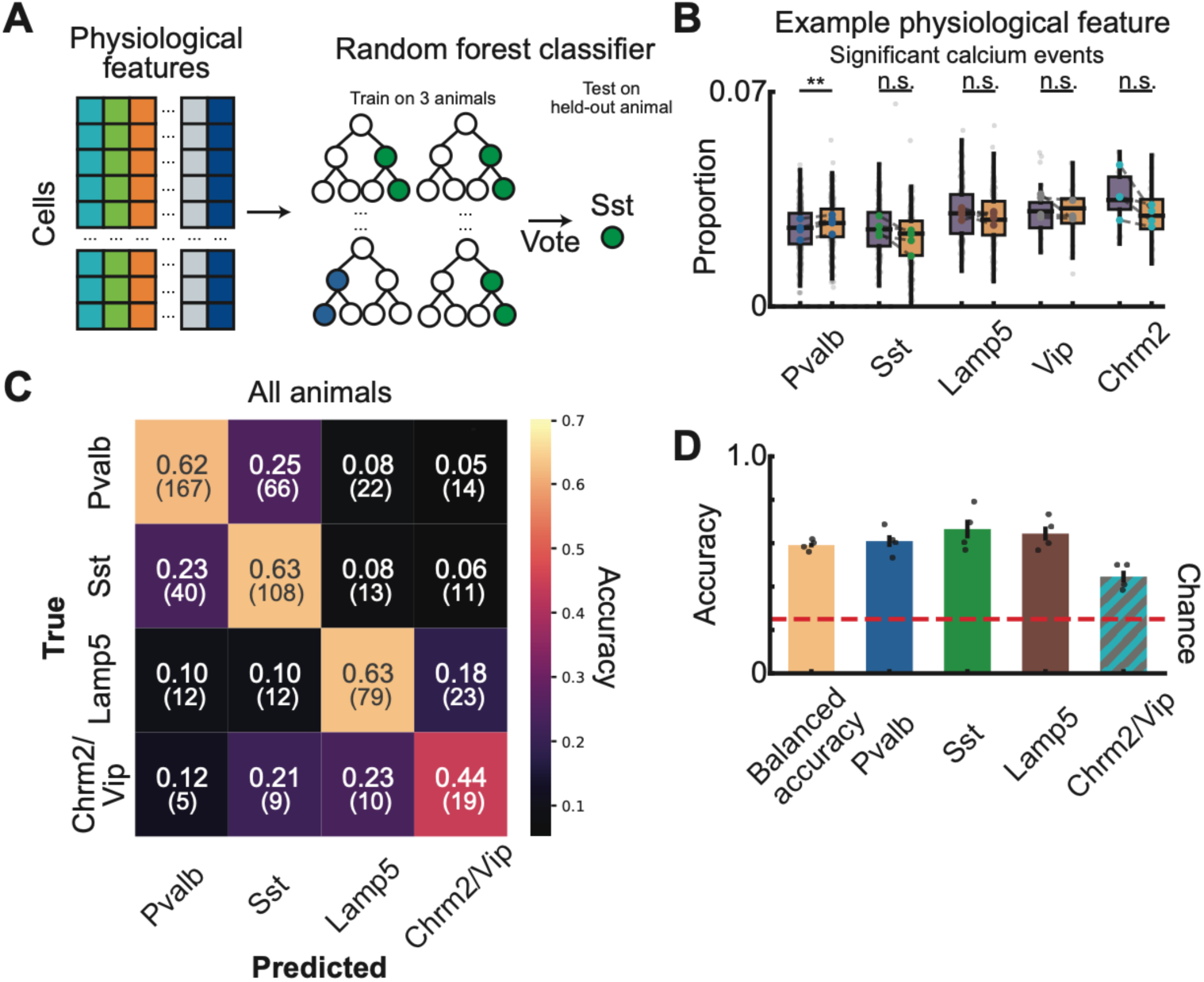
Physiology-based classification of CA1 IN subclasses. (**A**) Overview of the classification pipeline. A random forest classifier was trained using physiological features extracted from calcium traces to classify CA1 INs by subclass identity. For each fold, the classifier was trained on 3 animals and tested on 1 held-out animal (leave-one-animal-out cross-validation; see Methods). (**B**) An example physiological feature used in the classifier. The proportion of significant calcium events (defined as events exceeding 2 S.D. above the mean), calculated in familiar and novel contexts. (**C**) Confusion matrix pooled across all animals. Numbers indicate the fraction of cells in each true class assigned to each predicted class. Cell counts are in parentheses. (**D**) Overall balanced accuracy and subclass-wise accuracy across animals. Dots indicate individual animals and bars indicate mean accuracy. The red dashed line indicates the chance level (0.25). All accuracies were above chance (overall balanced accuracy: 0.59 ± 0.012 (mean ± SEM), t(3)=27.43, p=0.0001; *Pvalb*: 0.61 ± 0.031, t(3)=11.45, p=0.0014; *Sst*: 0.67 ± 0.050, t(3)=8.32, p=0.0036; *Lamp5*: 0.64 ± 0.037, t(3)=10.54, p=0.0018; *Chrm2*/*Vip*: 0.45 ± 0.031, t(3)=6.28, p=0.0082; n=4 animals for all comparisons; one-sample, two-sided t-test against chance).

**Table S1.**
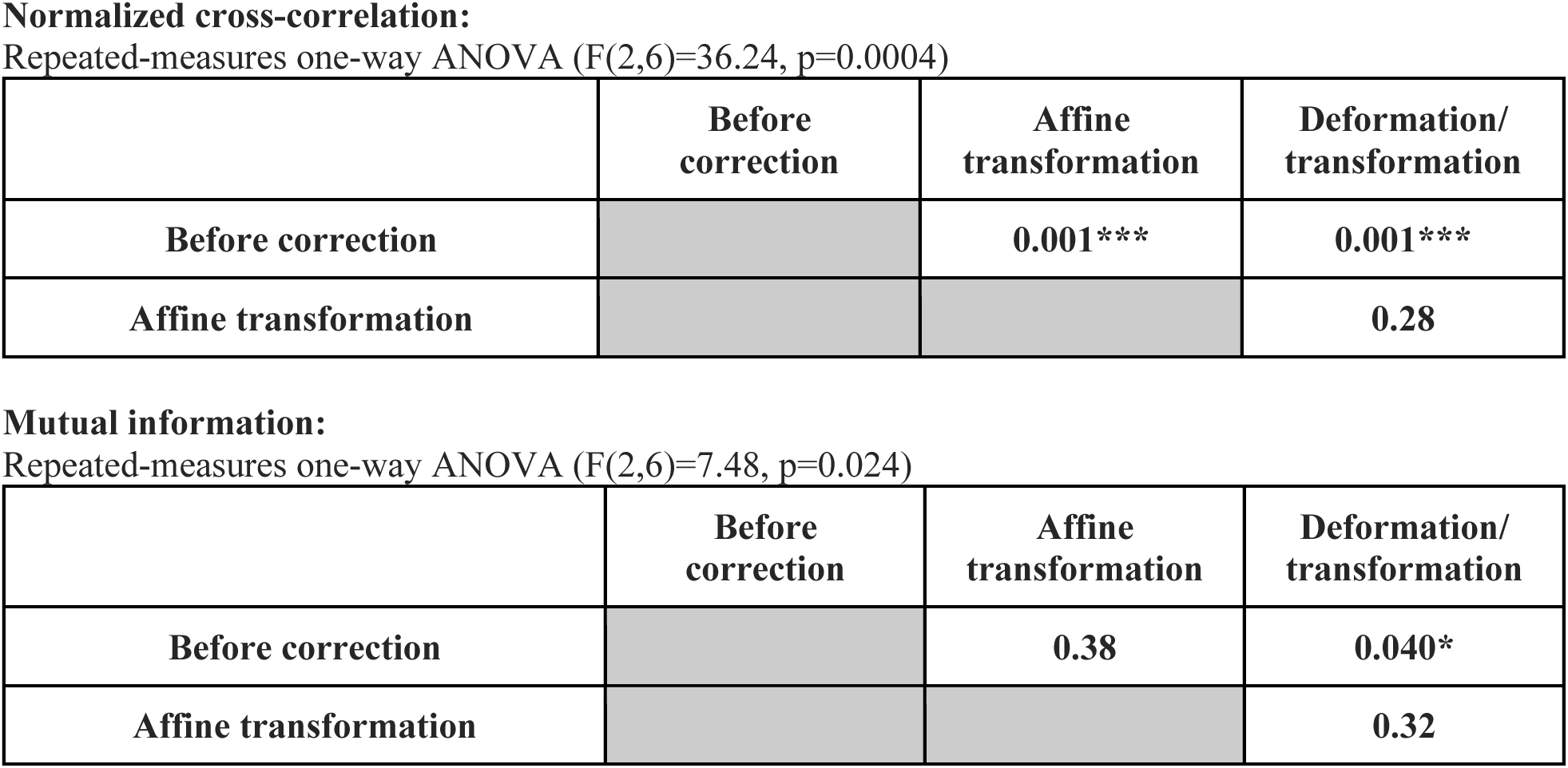
Statistics for registration metrics.

|  | Before correction | Affine transformation | Deformation/ transformation |
| --- | --- | --- | --- |
| Before correction |  | 0.001*** | 0.001*** |
| Affine transformation |  |  | 0.28 |

**Mutual information:** Repeated-measures one-way ANOVA (F(2,6)=7.48, p=0.024)
|  | Before correction | Affine transformation | Deformation/ transformation |
| --- | --- | --- | --- |
| Before correction |  | 0.38 | 0.040* |
| Affine transformation |  |  | 0.32 |

**Table S2.** Laminar distribution of registered CA1 INs by subclass.

|  | *Pvalb* | *Sst* | *Lamp5* | *Vip* | *Chrm2* | **Total** |
| --- | --- | --- | --- | --- | --- | --- |
| *Oriens* | 31.2% (116) | 45.2% (168) | 16.1% (60) | 2.2% (8) | 5.4% (20) | 372 |
| *Pyramidale* | 64.1% (150) | 1.7% (4) | 27.8% (65) | 6.0% (14) | 0.4% (1) | 234 |
| *Radiatum* | 75.0% (3) | - | 25.0% (1) | - | - | 4 |

**Table S3.**
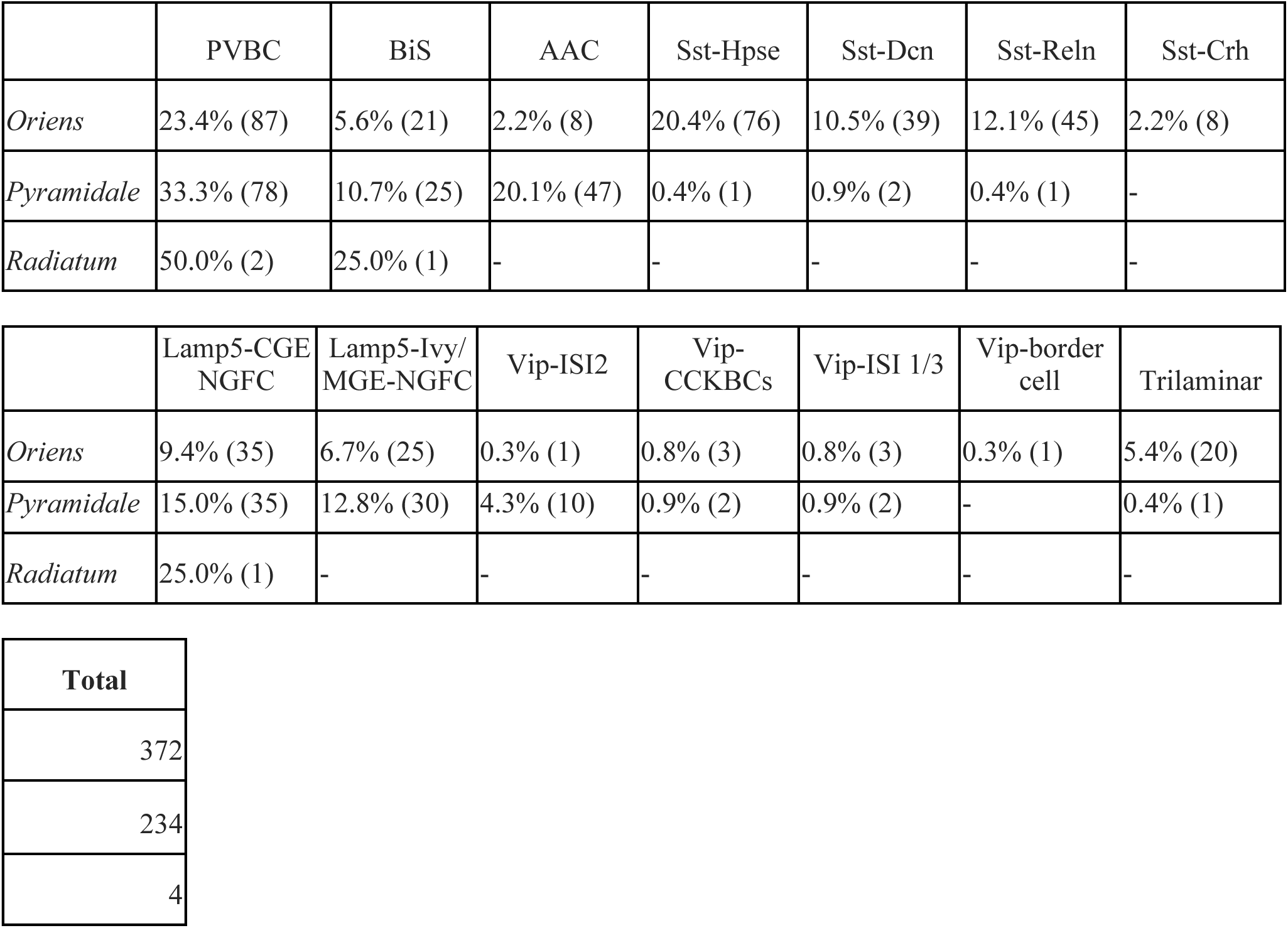
Laminar distribution of registered CA1 INs by type.

**Table S4.** Statistics for correlations between tPC2 and physiological features.

**Subclass**
| **metric** | **Rho**<br>**(subclass)** | **Uncorrected**<br>**P-value**<br>**(subclass)** | **FRD-**<br>**adjusted p-value**<br>**(subclass)** |
| --- | --- | --- | --- |
| Velocity modulation (familiar) | **-0.81** | **0.00002** | **0.00018** |
| Velocity modulation (novel) | **-0.68** | **0.00103** | **0.00464** |
| Rate map correlation (familiar vs. novel) | -0.36 | 0.11746 | 0.13214 |
| Change in sparsity (familiar vs. novel) | -0.36 | 0.11472 | 0.13214 |
| Reward modulation index (familiar) | **-0.57** | **0.00899** | **0.01618** |
| Reward modulation index (novel) | **-0.66** | **0.00162** | **0.00486** |
| Reward selectivity index (familiar) | **-0.63** | **0.00309** | **0.00695** |
| Reward selectivity index (novel) | -0.34 | 0.14703 | 0.14703 |
| Change in mean F/F | **0.51** | **0.02122** | **0.03183** |

**Type**
| **metric** | **Rho**<br>**(type)** | **Uncorrected**<br>**P-value (type)** | **FRD-**<br>**adjusted p-value**<br>**(type)** |
| --- | --- | --- | --- |
| Velocity modulation (familiar) | **-0.64** | **8.83e-07** | **7.94e-06** |
| Velocity modulation (novel) | **-0.52** | **0.00015** | **0.00045** |
| Rate map correlation (familiar vs. novel) | -0.28 | 0.05852 | 0.07524 |
| Change in sparsity (familiar vs. novel) | -0.18 | 0.20841 | 0.23446 |
| Reward modulation index (familiar) | **-0.37** | **0.00940** | **0.01410** |
| Reward modulation index (novel) | **-0.50** | **0.00032** | **0.00072** |
| Reward selectivity index (familiar) | **-0.56** | **0.00004** | **0.00018** |
| Reward selectivity index (novel) | -0.15 | 0.31881 | 0.31881 |
| Change in mean F/F | **0.48** | **0.00050** | **0.00090** |

**Table S5.** Features used in the physiology-based classifier.

| **Category** | **Variables** |
| --- | --- |
| **Space related** | Sparsity in the familiar context (F); sparsity in the novel context (N); rate-map correlation between F and N; rate-map correlation between F and N within individual blocks (block 1, laps 1–20; block 2, laps 21–40; block 3, laps 41–60); split-half reliability in F; split-half reliability in N |
| **Reward related** | Reward modulation index (RMI) in F; reward modulation index (RMI) in N; reward selectivity index (RSI) in F; reward selectivity index (RSI) in N; RMI for the first reward location in F; RMI for the second reward location in F; RMI for the first reward location in N; RMI for the second reward location in N |
| **Velocity related** | Velocity modulation in F; velocity modulation in N; velocity modulation during the familiar inter-trial interval (ITI); velocity modulation during the novel inter-trial interval (ITI) |
| **Activity level related** | Peak $\Delta F/F$ in F; peak $\Delta F/F$ in N; mean $\Delta F/F$ in F; mean $\Delta F/F$ in N; mean $\Delta F/F$ during the familiar ITI; mean $\Delta F/F$ during the novel ITI; proportion of significant calcium events in F; proportion of significant calcium events in N; proportion of significant calcium events during the familiar ITI; proportion of significant calcium events during the novel ITI |

**Table S6.**
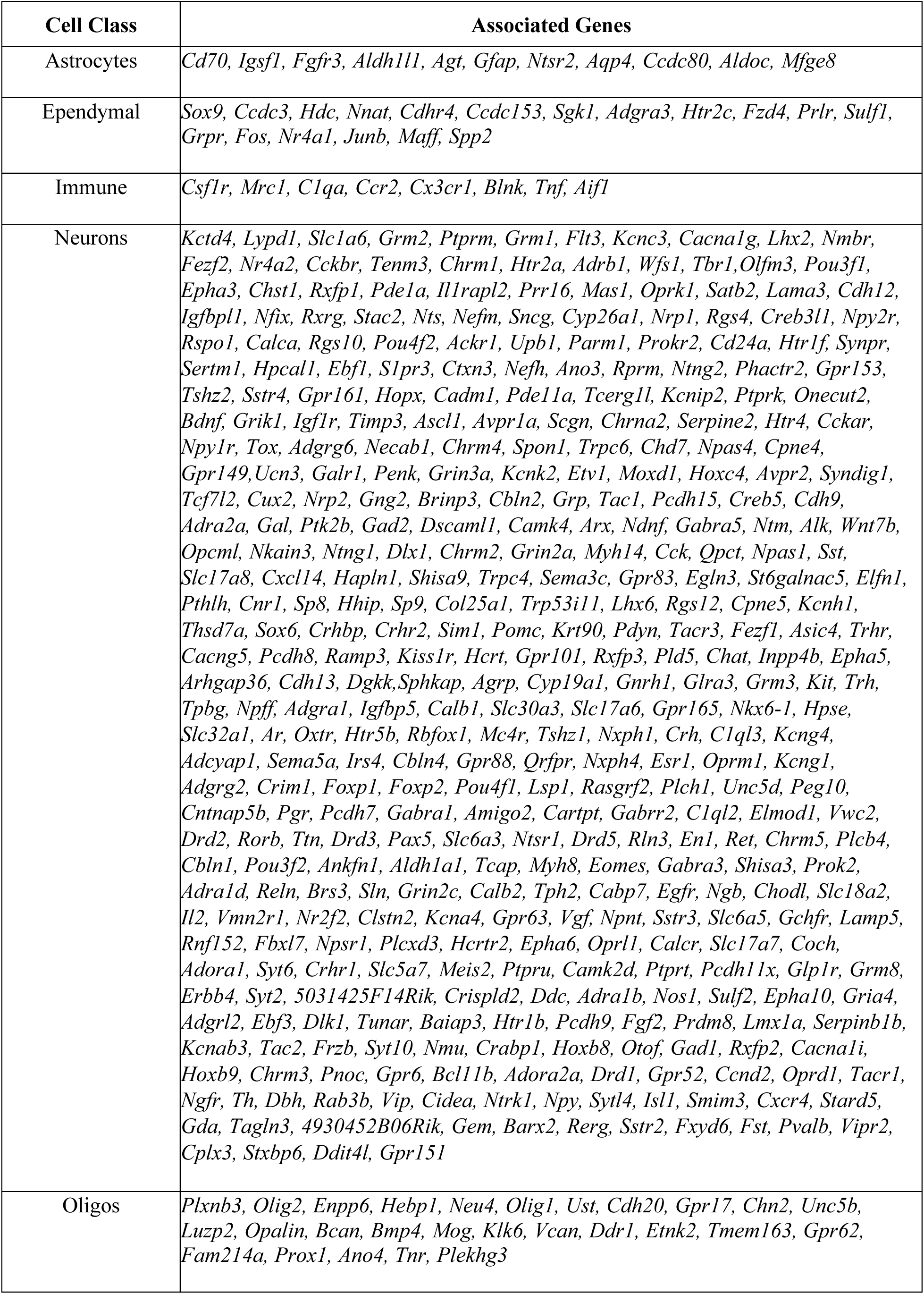
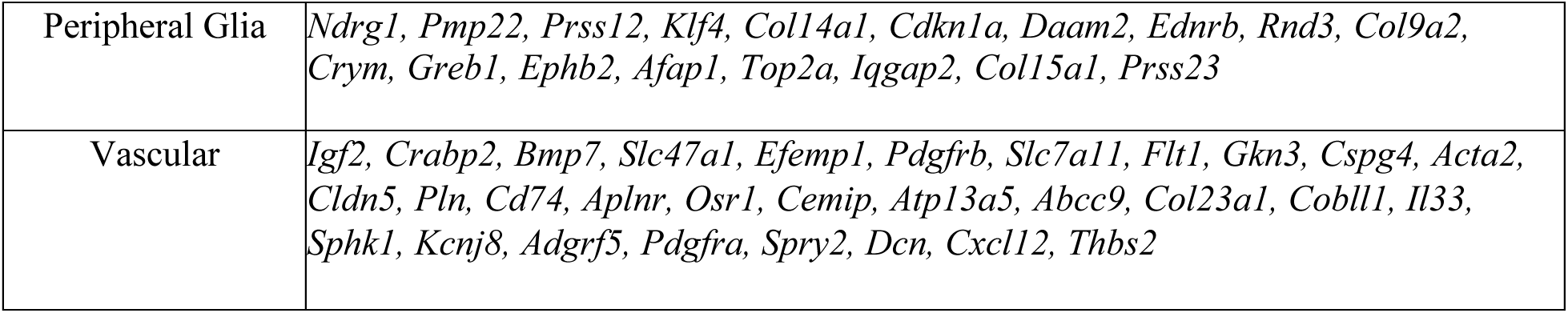
MERSCOPE PanNeuro cell type mouse panel (500 genes)

**Extended Data Figure 1.**
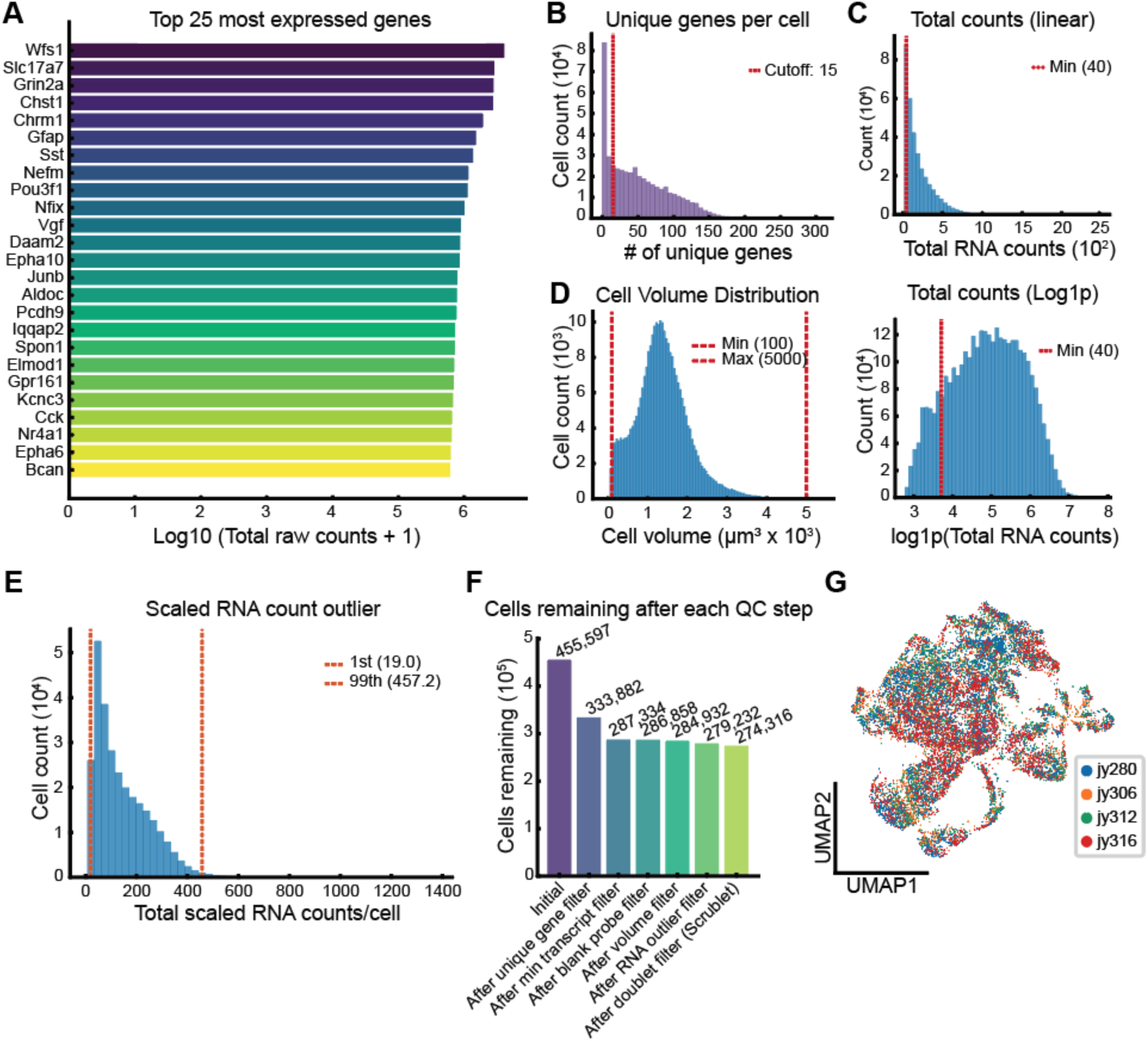
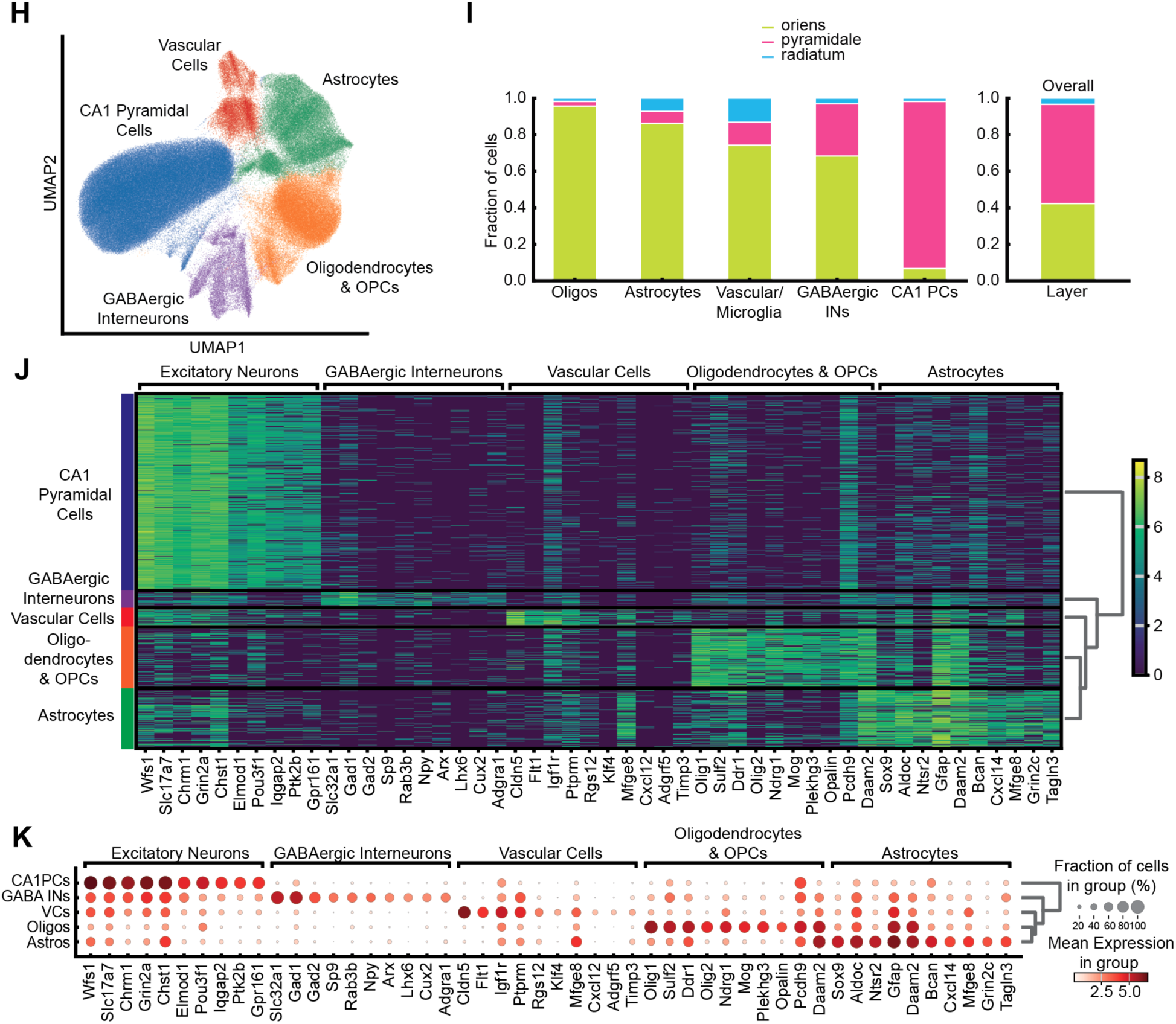
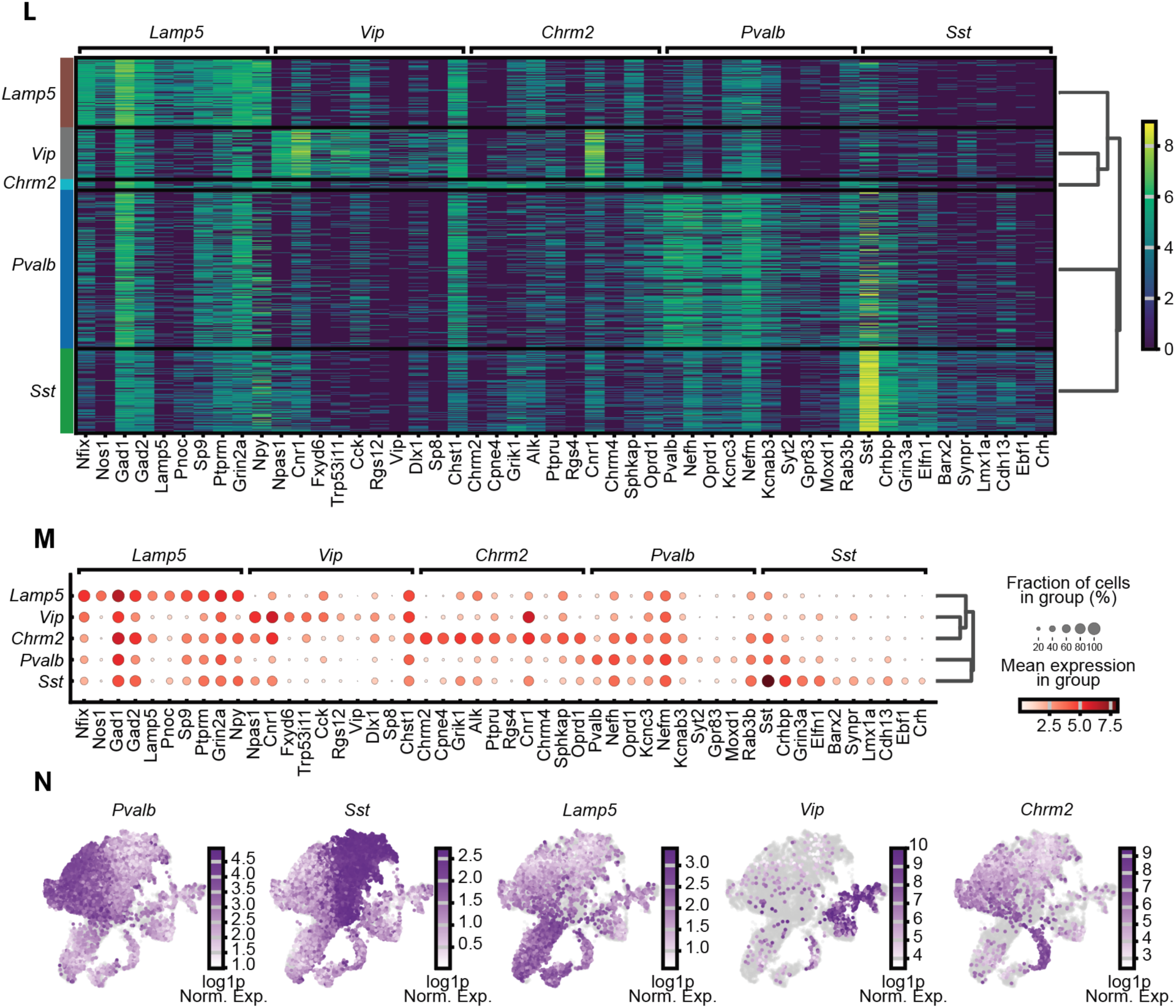
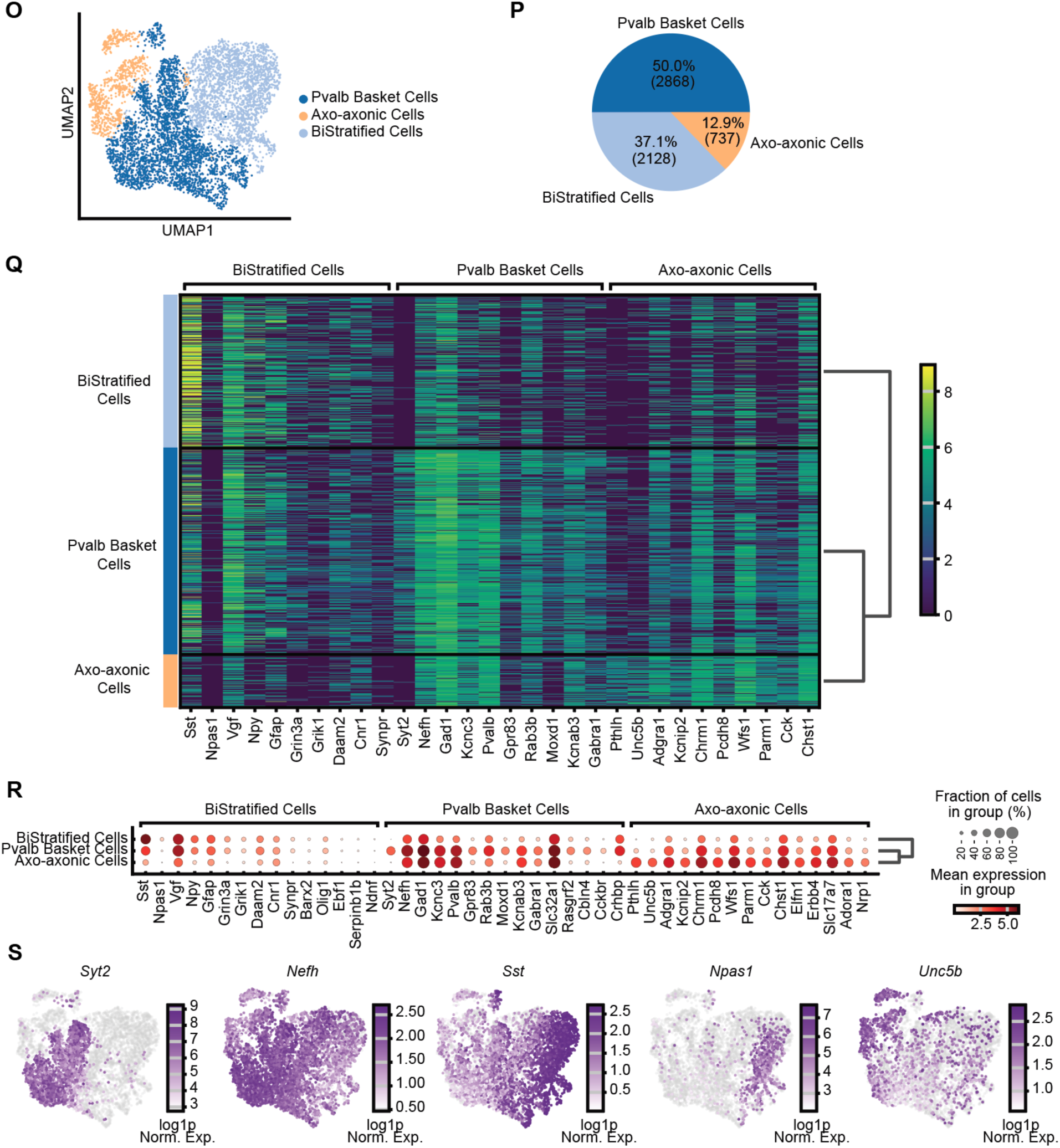
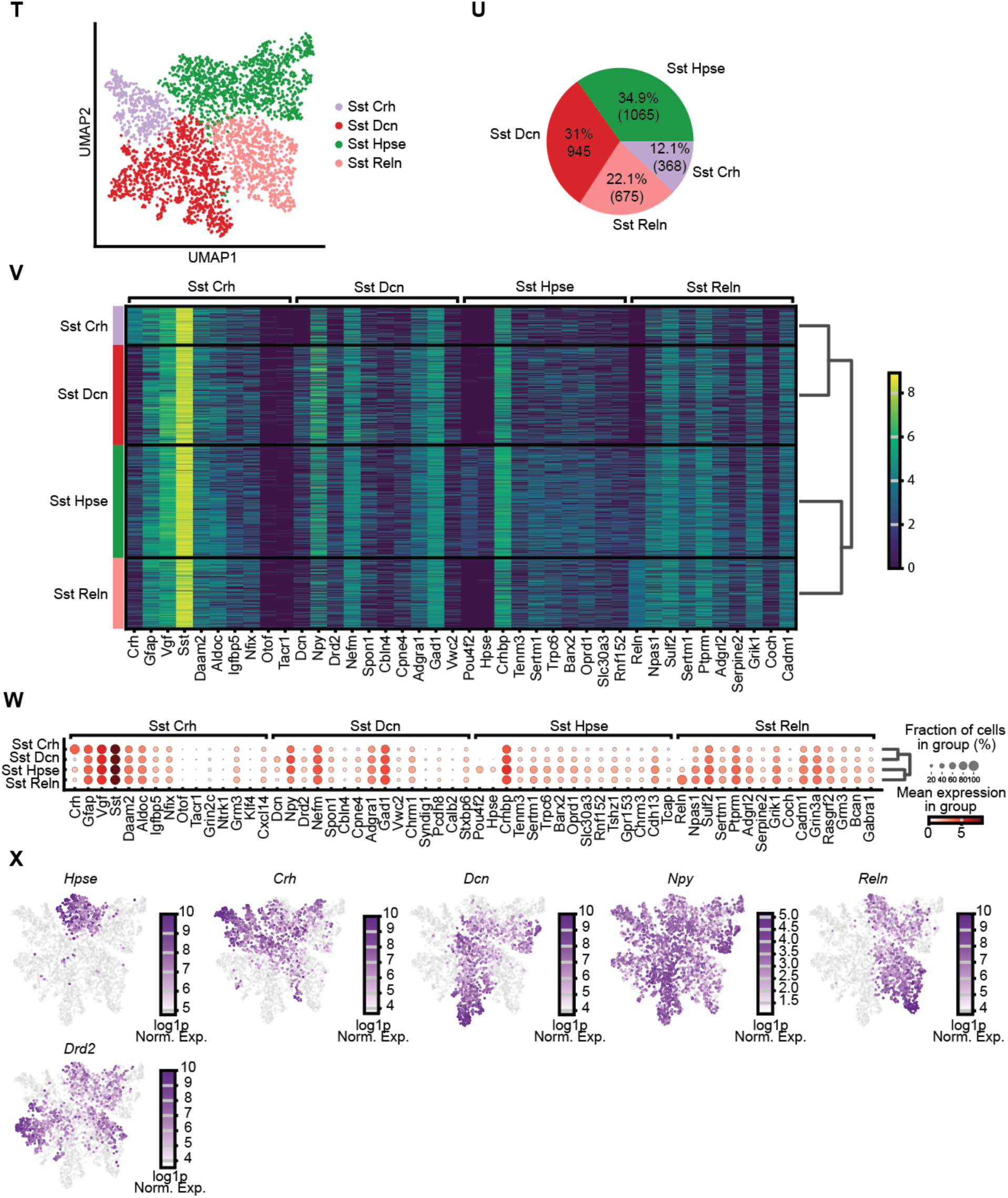
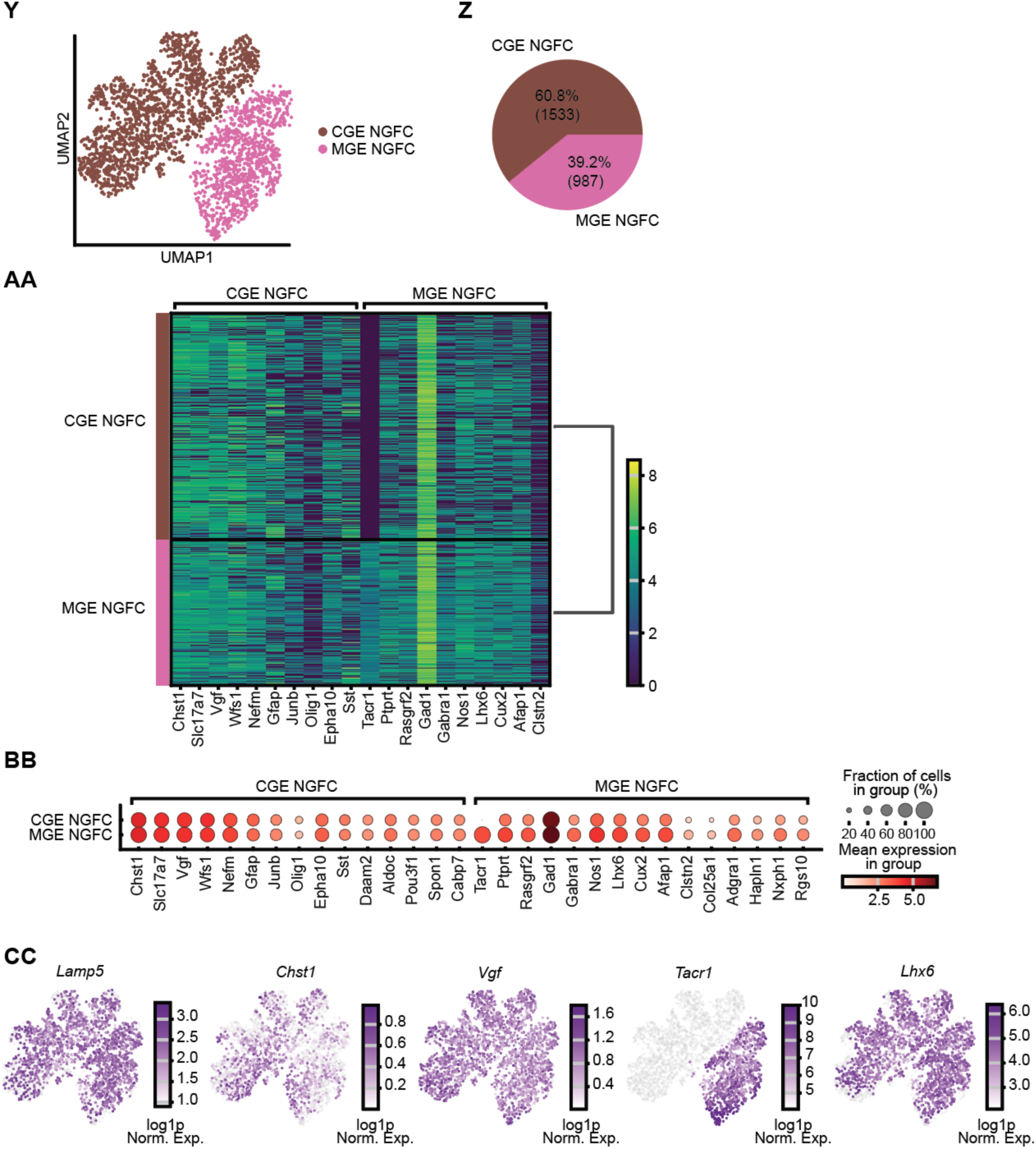
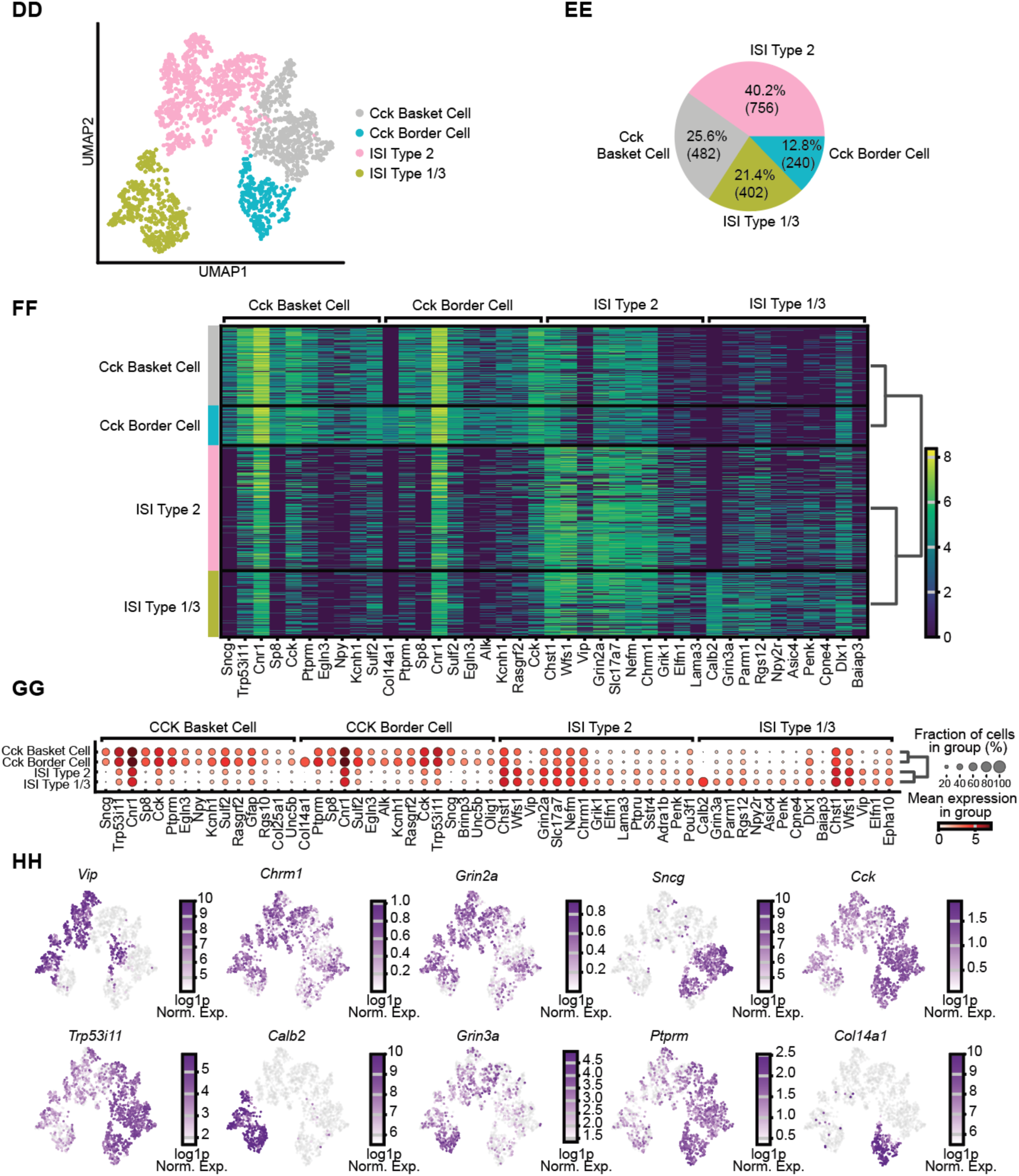
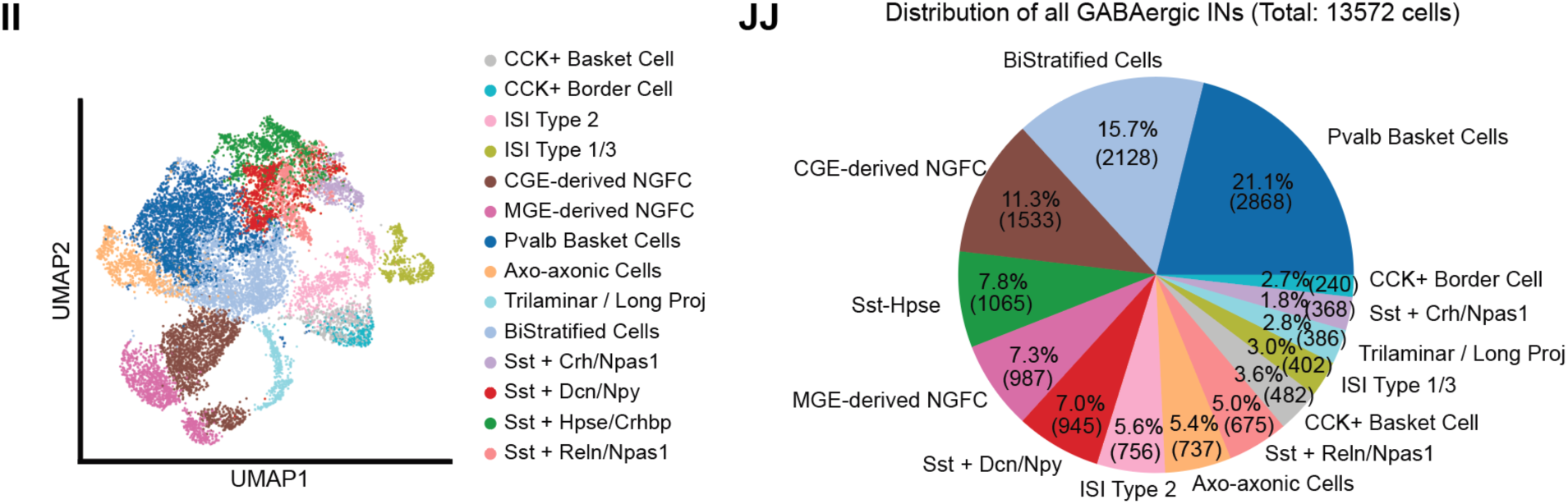
Unsupervised clustering pipeline and characterization for transcriptomically defined GABAergic cell types. (**A**) Top 25 highest expressed genes across the entire hippocampal dataset. (**B**) Distribution of the number of unique genes per cell. Red line depicts quality control cutoff (15 genes). (**C**) Distribution of total transcript counts per cell, visualized linearly (above) and log1p (below) of total RNA counts. Red line depicts minimum quality control cutoff (40 transcript counts). (**D**) Distribution of cell volumes across all cells. Red lines depict minimum (10 μm^3^ × 10^3^) and maximum (5000 μm^3^ × 10^3^) cell volume quality control cutoffs. (**E**) Distribution of scaled RNA counts. Red lines depict 1^st^ and 99^th^ percentile quality control cutoffs. (**F**) Histogram depicting cells remaining after each subsequent segmentation quality control step. (**G**) UMAP of all remaining cells colored by animal, depicting no obvious batch effects between animals (n = 4 animals, n = 274,316 cells). (**H**) UMAP visualization of 5 cell classes after unsupervised clustering. (**I**) Histogram depicting CA1 layer distribution of major cell classes. (**J**) Class level hierarchical heatmap depicting gene expression counts per cell of top 10 differentially expressed genes per class. (**K**) Dotplot depicting fraction of cells and mean expression of the top 10 differentially expressed genes per class. (**L**) GABAergic Interneuron subclass level hierarchical heatmap depicting gene expression counts per cell of top 10 differentially expressed genes per subclass. (**M**) Dotplot depicting fraction of cells and mean expression of the top 10 differentially expressed genes per GABAergic interneuron subclass. (**N**) UMAPs depicting subclass marker gene expression across all GABAergic interneurons. (**O**) UMAP depicts reclustering results of the *Pvalb* subclass, identifying 3 clusters: Pvalb Basket Cells, Axo-axonic cells, and BiStratified Cells (n = 4 animals, n = 5,733 cells). (**P**) Pie chart depicting cell cluster proportion and quantities within the *Pvalb* subclass. (**Q**) *Pvalb* subclass level hierarchical heatmap depicting gene expression counts per cell of top 10 differentially expressed genes per cluster. (**R**) Dotplot depicting fraction of cells and mean expression of the top 10 differentially expressed genes per *Pvalb* cluster. (**S**) UMAPs depicting cluster marker gene expression across *Pvalb* clusters. (**T**) UMAP depicts reclustering results of the *Sst* subclass, identifying 4 clusters: Sst Crh, Sst Dcn, Sst Hpse, and Sst Reln (n = 4 animals, n = 2,685 cells). (**U**) Pie chart depicting cell cluster proportion and quantities within the *Sst* subclass. (**V**) *Sst* subclass level hierarchical heatmap depicting gene expression counts per cell of top 10 differentially expressed genes per cluster. (**W**) Dotplot depicting fraction of cells and mean expression of the top 10 differentially expressed genes per *Sst* cluster. (**X**) UMAPs depicting cluster marker gene expression across all *Sst* clusters. (**Y**) UMAP depicting reclustering results of the *Lamp5* subclass, identifying 2 clusters: medial ganglionic eminence (MGE)-derived neurogliaform cells (NGFC)s, and caudal ganglionic eminence (CGE)-derived NGFCs (n = 4 animals, n = 2,520 cells). (**Z**) Pie chart depicting cell cluster proportion and quantities within the *Lamp5* subclass. (**AA**) *Lamp5* subclass level hierarchical heatmap depicting gene expression counts per cell of top 10 differentially expressed genes per cluster. (**BB**) Dotplot depicting fraction of cells and mean expression of the top 15 differentially expressed genes per *Lamp5* cluster. (**CC**) UMAPs depicting cluster marker gene expression across all *Lamp5* clusters. (**DD**) UMAP depicts reclustering results of the *Vip* subclass, identifying 4 clusters: Cck Basket Cells, Cck Border Cells, Interneuron Selective Interneuron (ISI) Type 2, and ISI Type 1/3 (n = 4 animals, n = 1,880 cells). (**EE**) Pie chart depicting cell cluster proportion and quantities within the *Vip* subclass. (**FF**) *Vip* subclass level hierarchical heatmap depicting gene expression counts per cell of top 10 differentially expressed genes per cluster. (**GG**) Dotplot depicting fraction of cells and mean expression of the top 15 differentially expressed genes per *Vip* cluster. (**HH**) UMAPs depicting cluster marker gene expression across all *Vip* clusters. (**II**) UMAP mapping cluster-level results of all GABAergic interneuron clusters, depicting all 14 clusters. (**JJ**) Pie chart depicting cell cluster proportion and quantities of all 14 GABAergic interneuron clusters (13,572 cells total). CA1PCs: CA1 pyramidal cells. GABA INs: GABAergic inhibitory interneurons. VCs: Vascular cells. Oligos: Oligodendrocytes and oligodendrocyte progenitors. Astros: Astrocytes. MGE: Medial ganglionic eminence. NGFCs: Neurogliaform cells. CGE: Caudal ganglionic eminence. ISI: Interneuron selective interneuron.

**Movie S1. Cross-modal registration overview.** Animation illustrating the registration pipeline used to track CA1 interneurons from *in vivo* two-photon calcium imaging to *ex vivo* confocal imaging and MERSCOPE spatial transcriptomics, demonstrating reliable alignment across imaging modalities.

## Notes

### Competing Interest Statement

The authors have declared no competing interest.

